# Retinoic acid resolves the conflict between X-chromosome inactivation and pluripotency program in female cleavage-stage embryos

**DOI:** 10.64898/2026.04.12.717983

**Authors:** Qianying Yang, Lei An, Wenjing Wang, Shuai Zhang, Lizhu Ma, Qingji Lyu, Yuan Yue, Hong Deng, Chao Zhang, Xiao Hu, Juan Liu, Meiqiang Chu, Yawen Tang, Xiaodong Wang, Zhenni Zhang, Wei Fu, Jun Wu, Jianhui Tian

## Abstract

The tight coupling of X-chromosome inactivation (XCI) and pluripotency is a paradigm of holistic developmental regulation, and their inverse correlation enables XCI establishment during implantation. In contrast, the co-establishment of imprinted XCI (iXCI) and pluripotency program in female preimplantation embryos challenges the holistic pattern, but how embryos coordinate the two remains unknown. Here, we find that *Nanog* during the cleavage stage strongly represses the co-upregulated *Rnf12*-*Xist* axis, the triggering signaling of iXCI, creating a conflict that threatens iXCI. We identify retinoic acid (RA) as the coordinator that resolves the conflict and safeguards iXCI. RA activates its receptor RARG to evict NANOG from *Rnf12* 5’region, avoiding NANOG’s threat. *In vivo*, either maternal vitamin A deficiency or embryonic RA synthesis defects results in impaired iXCI and female-biased embryonic lethality. Thus, we highlight RA as a master regulator to holistically balance iXCI and pluripotency during cleavage stages, thereby ensuring proper female embryogenesis.

## INTRODUCTION

X-chromosome inactivation (XCI) is a female-specific process essential for balancing gene expression between the sexes in mammals ^1,2^. Mouse embryos undergo two waves of XCI: imprinted XCI (iXCI) initiates during early preimplantation dependent on the paternally expressed long non-coding RNA *Xist*, while random XCI (rXCI) occurs during peri-implantation via randomly upregulated monoallelic *Xist* ^3–5^. It has been well-established that pluripotency factors strongly repress *Xist* in pluripotent cells thus rXCI only initiates upon differentiation, and their tight coupling is thought to be a classical paradigm of holistic developmental regulation ^6–8^.

The acquisition of pluripotency in preimplantation embryos is the prerequisite for establishing developmental potential ^9^. However, in sharp contrast to the loss of pluripotency that derepresses rXCI initiation, two conflicting yet essential processes must co-establish simultaneously during the early cleavage stage (Figures 1A and S1A), but how female preimplantation embryos coordinate the two remains unknown. In particular, our previous study in *in vitro* fertilized (IVF) embryos ^10^, as well as a genetic model with enforced *Nanog* ^11^, suggest that *Nanog*, which occupies a central position in preimplantation pluripotency acquisition ^9^, appears to impede *Xist* upregulation and thus threatens iXCI, highlighting the need to understand how preimplantation embryos resolve the conflict between iXCI and pluripotency.

**Figure 1.**
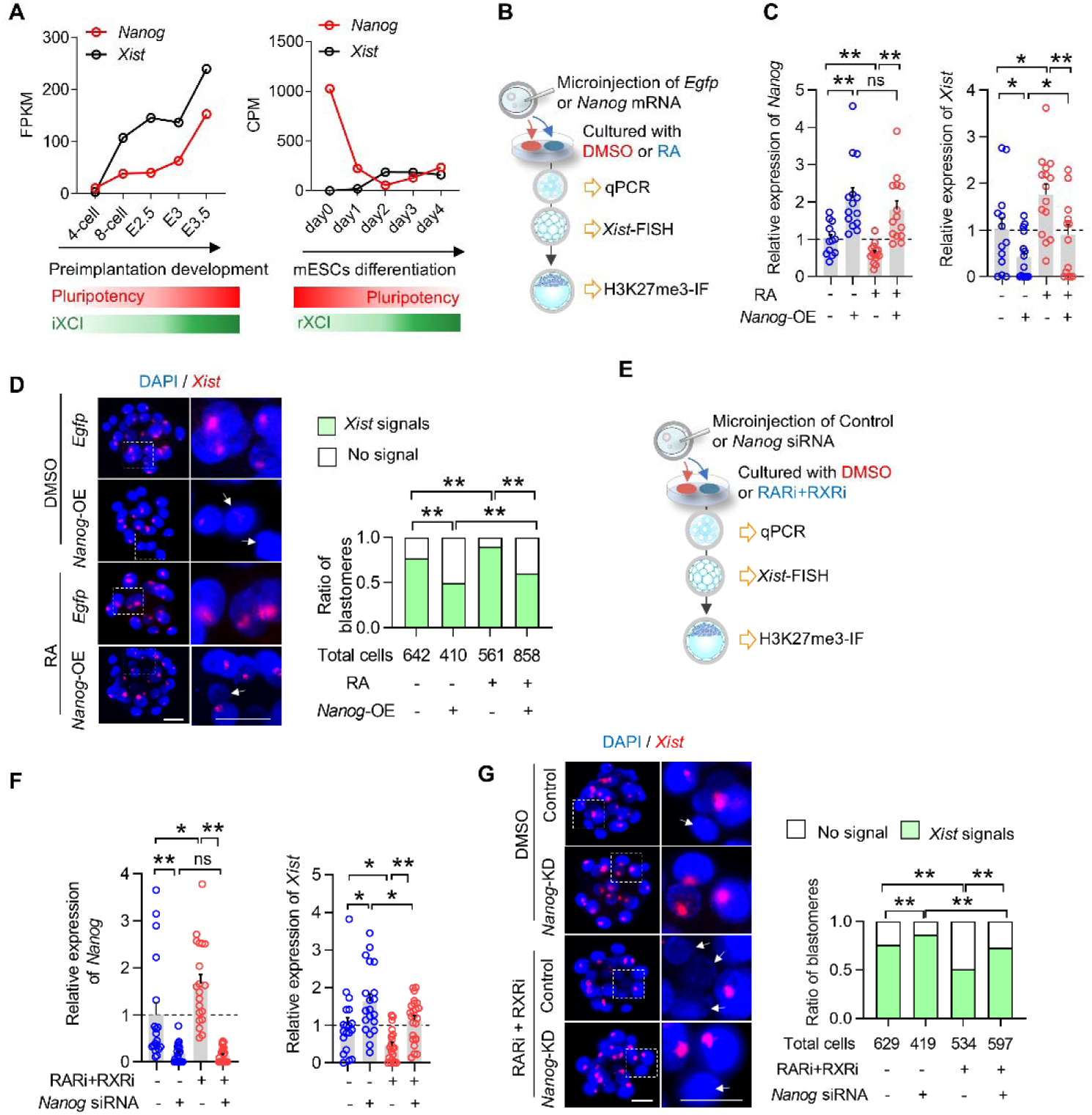
NANOG efficiently represses iXCI establishment during early cleavage stage and RA antagonizes NANOG’s repression effect. **(A)** Expression dynamics of *Xist* and *Nanog* during preimplantation development and ES cell differentiation. (**B**) Schematic diagram of the experimental workflow. *Nanog* was overexpressed by microinjecting *Nanog* mRNA into zygotes in the presence or absence of exogenous RA, hallmarks of XCI establishment, including *Xist* upregulation, *Xist* RNA coating, H3K27me3 enrichment were detected at the 8-cell, morula and blastocyst stages respectively. (**C**) Relative expression levels of *Nanog* and *Xist* in 8-cell female embryos subjected to RA supplementation or *Nanog* overexpression, either alone or in combination. Each dot represents a single female 8-cell embryo. (**D**) *Xist-*FISH analysis in female morulae. The right column shows a higher magnification of the boxed region. White arrows indicate blastomeres without the *Xist* RNA domain. Blue arrows indicate blastomeres with *Xist* RNA domain. Right panel: the percentage of *Xist*-positive and -negative blastomeres to total blastomeres in female morulae. (**E**) Schematic diagram of the experimental workflow. *Rnf12* knockdown was performed by microinjecting *Rnf12* siRNA into zygotes in the presence or absence of RAR/RXR pan-inhibitors. *Xist* upregulation, *Xist* RNA coating, H3K27me3 enrichment were detected at the 8-cell, morula and blastocyst stages respectively. **(F)** Relative expression levels of *Nanog* and *Xist* in female 8-cell embryos subjected to RAR/RXR pan-inhibition or *Nanog* knockdown, either alone or in combination. Each dot represents a single female 8-cell embryo. **(G)** *Xist-*FISH analysis in female morulae from different intersections and maternal dietary conditions. White arrows indicate blastomeres without the *Xist* RNA domain. Right panel: the percentage of *Xist*-positive and -negative blastomeres to total blastomeres in female morulae. data are mean ± s.e.m and two-tailed Student’s t-tests were used to determine statistical significance. \**p* < 0.05, \*\**p* < 0.01, ns, not significant. Scale bar = 25 μm.

Retinoic acid (RA) has been used for decades to induce ES cell differentiation and accompanying rXCI. RA also finetunes pluripotency factors to orchestrate the pluripotency program ^12,13^. Our previous study has revealed that exogenous RA supplementation in IVF culture medium not only restrains excessive upregulation of *Nanog*, but also rescues iXCI upregulation ^10^. Given RA is present in oviductal fluid, which is critical for supporting early development, we ask whether RA is the physiological coordinator that orchestrates simultaneous upregulation of *Xist* and *Nanog*, thus holistically ensuring concurrent iXCI establishment and pluripotency program.

Here, we identify RA as the critical coordinator that orchestrates simultaneous iXCI and pluripotency program during preimplantation development. In the absence of RA signaling, *Nanog* strongly represses of the *Rnf12-Xis*t axis, the triggering signaling of iXCI, thus raising a conflict that impairs iXCI. Within a multifaceted circuitry, RA resolves the conflict by activating its receptor RARG to evict NANOG from the regulatory elements of *Rnf12* 5’region, thus avoiding NANOG’s repression of iXCI. Our findings not only highlight the role of RA as a master regulator orchestrating a critical conflict-resolution mechanism to ensure proper female embryogenesis, but also establish a mechanistic link between a vital nutrient and offspring sex ratio, elucidating how VA deficiency skews sex ratio at a population level.

## RESULTS

### NANOG represses iXCI establishment at the early cleavage stage and RA antagonizes this repression

Given the simultaneous co-upregulation of *Nanog* and *Xist* during preimplantation embryos, as well as the excessively upregulated *Nanog* and impaired iXCI observed in IVF embryos ^10^, we initially asked if NANOG represses *Xist* and is disruptive to iXCI establishment, thus raising a conflict. To test this, we overexpressed *Nanog* in preimplantation embryos during iXCI establishment (Figure. 1B and Figure. S1B). The excessively upregulated *Nanog* reduced *Xist* expression (Figure. 1C), and diminished *Xist* RNA coating (Figure. 1D and Figure. S1C) and caused a loss of H3K27me3 domains (Figure. S1D-E), the key markers for the initiation and establishment of the repressive chromatin state of the inactive X ^14^. Conversely, knocking down *Nanog* increased *Xist* expression (Figure. 1E-F and Figure. S1F) and improved iXCI status (Figure. 1G, Figure. S1G-J), demonstrating that *Nanog* serves as a repressor of *Xist* that threatens iXCI establishment.

Of note, NANOG’s repression can be significantly reversed by exogenous RA supplementation (Figure. 1C, D and Figure. S1C-E), while *Nanog* knockdown-induced *Xist* upregulation and increase in H3K27me3 domains were abolished by RA signaling blockage via pan-RAR/RXR inhibition (Figure. 1F-G, Figure. S1G-J). It should be also highlighted that in the absence of RA signaling, *Nanog* showed excessively upregulation while exogenous RA restrained *Nanog* (Figure. 1C, F). These results suggest that RA antagonizes NANOG’s repression to safeguards iXCI establishment in cleavage-stage embryos, thereby overcoming the developmental conflict imposed by *Nanog* upregulation and pluripotency program.

To directly confirm the critical role of RA in safeguarding iXCI, we took advantage of pan-RAR/RXR inhibition, by which nuclear RA receptor signaling can be completely blocked in cells (Figure. S2A) and cleavage-stage embryos (Figure. S2B). RA signaling blockage alone resulted in a severe loss of H3K27me3 (Figure. S2C-D) and *Xist* domains (Figure. S2E). Importantly, the impaired iXCI caused by cleavage-stage inhibition was irreversible, even when blastocysts were later transferred into the normal culture medium for outgrowth (Figure. S2F-H). Taken together, our data indicate that *Nanog* represses *Xist* during the cleavage stage and thus raises a conflict that is disruptive to iXCI in preimplantation embryos. RA antagonizes *Nanog*’s repression to safeguard *Xist* upregulation, thus ensures iXCI establishment.

### RA antagonizes *Nanog* and safeguards *Xist* upregulation via *Rnf12*

We next investigated how RA antagonizes NANOG and safeguards *Xist* upregulation to resolve the conflict between iXCI establishment and pluripotency program in cleavage-stage embryos. Since the results of single-embryo qRT-PCR analyses showed that RA doesn’t regulates *Xist* and *Nanog* until 8-cell stage (Figure. 1C, F and Figure. S3A), so we performed all subsequent transcriptional analyses in 8-cell embryos. Surprisingly, ATAC-seq and RAR/RXR ChIP-seq revealed that the potential binding region of nuclear retinoid receptors and NANOG on *Xist* intron was not accessible, suggesting that neither nuclear retinoid receptors nor NANOG can directly bind *Xist* intron during the iXCI establishment (Figure. 2A), as confirmed by iCUT&RUN-qPCR analyses of primarily expressed retinoid receptors (*Rarg* and *Rxra*) ^15^ during preimplantation development (Figure. 2B and Figure. S3B). These findings suggest that RA safeguards *Xist* upregulation and resolves the conflict via unknown mediator. Considering *Xist*’s sensitivity to RA signaling, we hypothesized the mediator may be also RA-responsive. Thus, we constructed a transcriptome map illustrating the embryonic response to either RA supplementation or retinoid receptor blockage at the 8-cell stage (Figure. 2C and Figure. S3C-D). The transcriptome analysis (Figure. S3E-G) and single-embryo RT-qPCR (Figure. 2D) further highlighted *Nanog*, but not other pluripotency factors, was highly sensitive to RA signaling.

**Figure 2.**
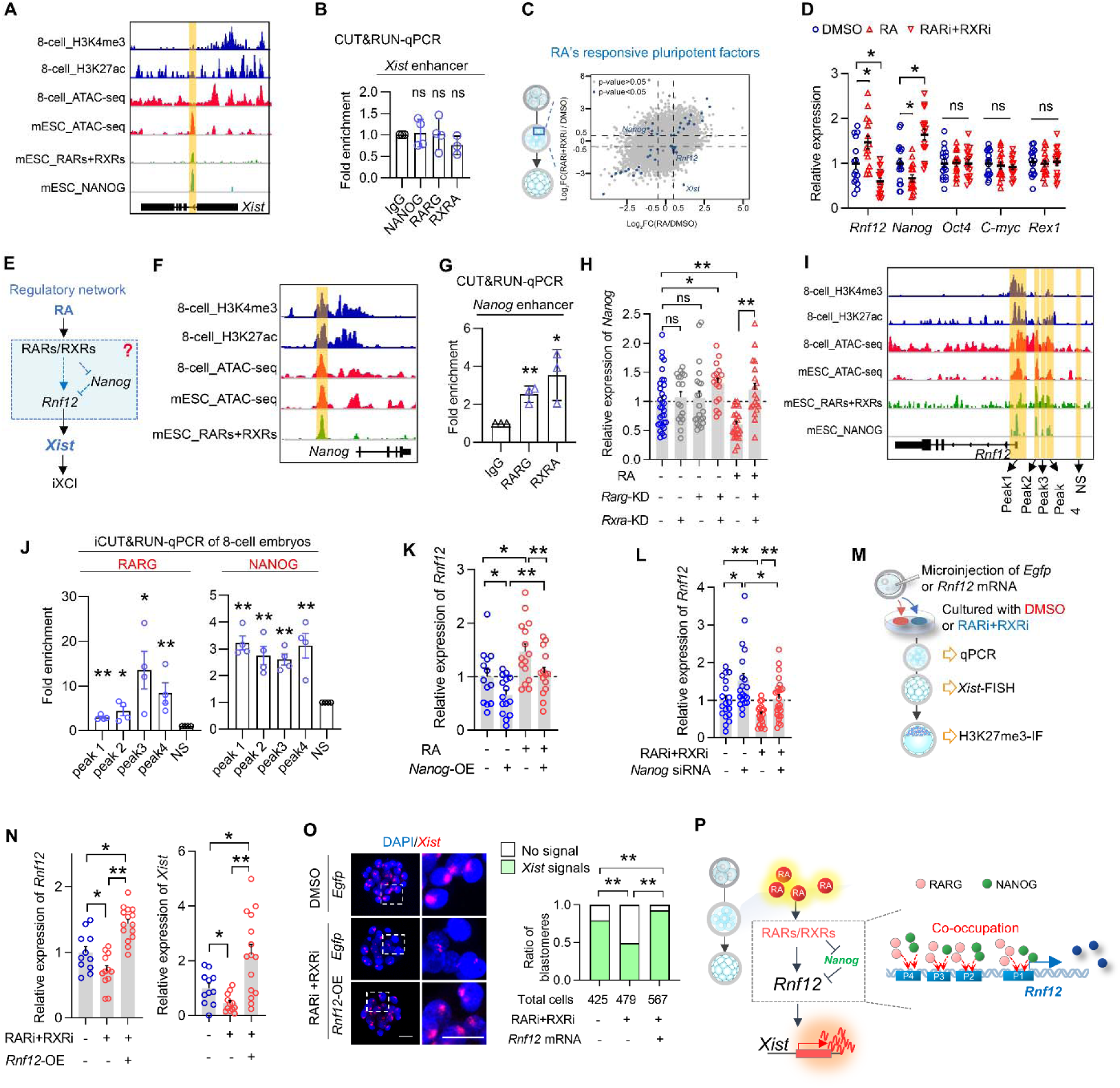
RA antagonizes *Nanog* and safeguards *Xist* upregulation via *Rnf12*. **(A)** Genome browser view of H3K4me3, H3K27ac, nuclear retinoid receptors (RARs+RXRs) and NANOG ChIP-seq and ATAC-seq in 8-cell embryos and ES cells around the *Xist* locus. The putative binding site of nuclear retinoid receptors based on their enrichment in mouse mESCs is highlighted. **(B)** iCUT&RUN-qPCR analysis of nuclear retinoid receptors and NANOG occupancy at their putative binding sites within regulatory regions of *Xist* in 8-cell embryos. Each dot represents a biological replicate (50 8-cell embryos per replicate). **(C)** Transcriptome map of the candidate mediator responding to both RA supplementation and RA signaling blockage. The blue dot indicates candidates that show significant bidirectional changes. Right panel: hypothesized model for RA regulatory mechanisms. **(D)** Relative expression levels of *Rnf12*, *Nanog*, *Oct4*, *C-myc*, *Rex1* in female 8-cell embryos exposed to exogenous RA or RAR/RXR pan-inhibitors. Each dot represents a single female 8-cell embryo. **(E)** Hypothesized model for RA regulatory mechanisms. **(F)** Genome browser view of H3K4me3, H3K27ac, RARs+RXRs ChIP-seq and ATAC-seq in 8-cell embryos and ES cells around the *Nanog* locus. The binding site on *Nanog* distal enhancer is highlighted. **(G)** iCUT&RUN-qPCR analysis of nuclear retinoid receptors occupancy at their putative binding sites within regulatory regions *Nanog* in 8-cell embryos. Each dot represents a biological replicate (50 8-cell embryos per replicate). **(H)** Relative expression levels of *Nanog* in female 8-cell embryos subjected to *Rarg* knockdown, *Rxra* knockdown, or RA supplementation, either alone or in combination. The results showed that RA regulates *Nanog* expression through RARG/RXRA. Each dot represents a single female 8-cell embryo. **(I)** Genome browser view of H3K4me3, H3K27ac, RARs+RXRs and NANOG ChIP-seq and ATAC-seq in 8-cell embryos and ES cells around the *Rnf12* locus. The putative binding sites (peak1-peak4) of nuclear retinoid receptors and NANOG are highlighted. NS, non-specific binding site. **(J)** iCUT&RUN-qPCR analysis of RARG and NANOG occupancy at their putative binding sites (peak1-peak4) within *Rnf12* 5’ region in 8-cell embryos. NS, non-specific binding site. Each dot represents a biological replicate (50 8-cell embryos per replicate). **(K)** Relative expression levels of *Rnf12* in female 8-cell embryos subjected to RA supplementation or *Nanog* overexpression, either alone or in combination. Each dot represents a single female 8-cell embryo. **(L)** Relative expression levels of *Rnf12* in female 8-cell embryos subjected to RAR/RXR pan-inhibition or *Nanog* knockdown, either alone or in combination. Each dot represents a single female 8-cell embryo. **(M)** Schematic diagram of the experimental workflow. *Rnf12* was overexpressed in the presence or absence of RAR/RXR pan-inhibitors, *Xist* upregulation, *Xist* RNA coating, H3K27me3 enrichment were detected at the 8-cell, morula and blastocyst stages respectively. **(N)** Relative expression levels of *Rnf12* and *Xist* in female 8-cell embryos exposed to RAR/RXR pan-inhibition alone or in combination with *Rnf12* overexpression. Each dot represents a single female 8-cell embryo. **(O)** *Xist-*FISH analysis in female morulae. Right panel: the percentage of *Xist*-positive and -negative blastomeres to total blastomeres in female morulae. **(P)** Schematic diagram of the model in which RA regulates iXCI. For n, Chi-square tests were used. For others, data are mean ± s.e.m and two-tailed Student’s t-tests were used to determine statistical significance. \**p* < 0.05, \*\**p* < 0.01, ns, not significant. Scale bar = 25 μm.

More importantly, among the known XCI regulators (Figure. S3E), *Rnf12*, an known essential activator for initiating *Xist* and iXCI ^4,6,7,16^, caught our attention. *Rnf12* not only responded sensitively to RA signaling (Figure. 2D and Figure. S3F-G), but also was identified as a direct target of NANOG in mESCs ^8^, suggesting that *Nanog* and *Rnf12* may construct the regulatory network that bridges RA signaling and *Xist* upregulation and resolves the conflict (Figure. 2E). In this network, nuclear retinoid receptors RARG and RXRA could directly bind to a distal enhancer ^17^ of *Nanog* (Figure. 2F-G) and synergistically restrained *Nanog* expression in a RA-responsive manner (Figure.2H and Figure. S3H). We next asked how *Rnf12* mediates the network. The integrated analyses of ATAC-seq and ChIP-seq revealed that both nuclear retinoid receptors and NANOG bind to the accessible regulatory regions of *Rnf12* in preimplantation embryos (Figure. 2I). Among the nuclear retinoid receptors expressed during preimplantation development (Figure. S3B), iCUT&RUN-qPCR assay showed that RARG binds to *Rnf12* 5’ region. More interestingly, promoter (peak1) and putative enhancer elements (peak2-4) within *Rnf12* 5’region were co-occupied by RARG and NANOG (Figure. 2J and Figure. S3I). These findings, together with the results that *Rnf12* was positively and negatively regulated by RA and NANOG respectively (Figure. 2K-L), suggest that *Rnf12*, as the direct target of both RARG and NANOG, serves as a hub gene in constructing the regulatory network responsible for resolving the conflict between the co-establishment of iXCI and pluripotency program (Figure. 2E).

To test the central role of *Rnf12*, we overexpressed it in RA signaling-blocked embryos (Figure. 2M and Figure. S4A). The restored *Xist* expression (Figure. 2N), *Xist* RNA coating (Figure. 2O and Figure. S4B-C) and H3K27me3 domains (Figure. S4D-F) confirmed our hypothesis. Thus, our data demonstrate that RA activates *Rnf12* via two parallel pathways: (1) direct transcriptional activation through RA-RARG binding to *Rnf12* promoter and enhancer regions; and (2) restraining excessive *Nanog* expression, which would otherwise upregulate *Rnf12* transcription. Additionally, RARG and NANOG co-occupy regulatory elements at *Rnf12* 5’ region, led us to ask their interaction on these regulatory elements may play a more direct role in resolving the conflict (Figure. 2P).

### RA activates RARG to evict NANOG from the 5’region of *Rnf12*

Because RARG and NANOG co-occupy the regulatory elements of *Rnf12* (Figure. 2I-J), and the absence of RA exacerbates this conflict (Figure. 1F-G, Figure. S1G-J), we hypothesized that RA-RARG coordinates the binding of NANOG on *Rnf12*’s regulatory elements to resolve the conflict. To test this, we examined whether: (1) RARG mediates RA’s role in *Rnf12* induction; (2) RARG competes with NANOG for binding to the regulatory elements of *Rnf12*; and (3) RA enhances RARG’s ability to evict NANOG from the *Rnf12* 5’region (Figure. 3A).

**Figure 3.**
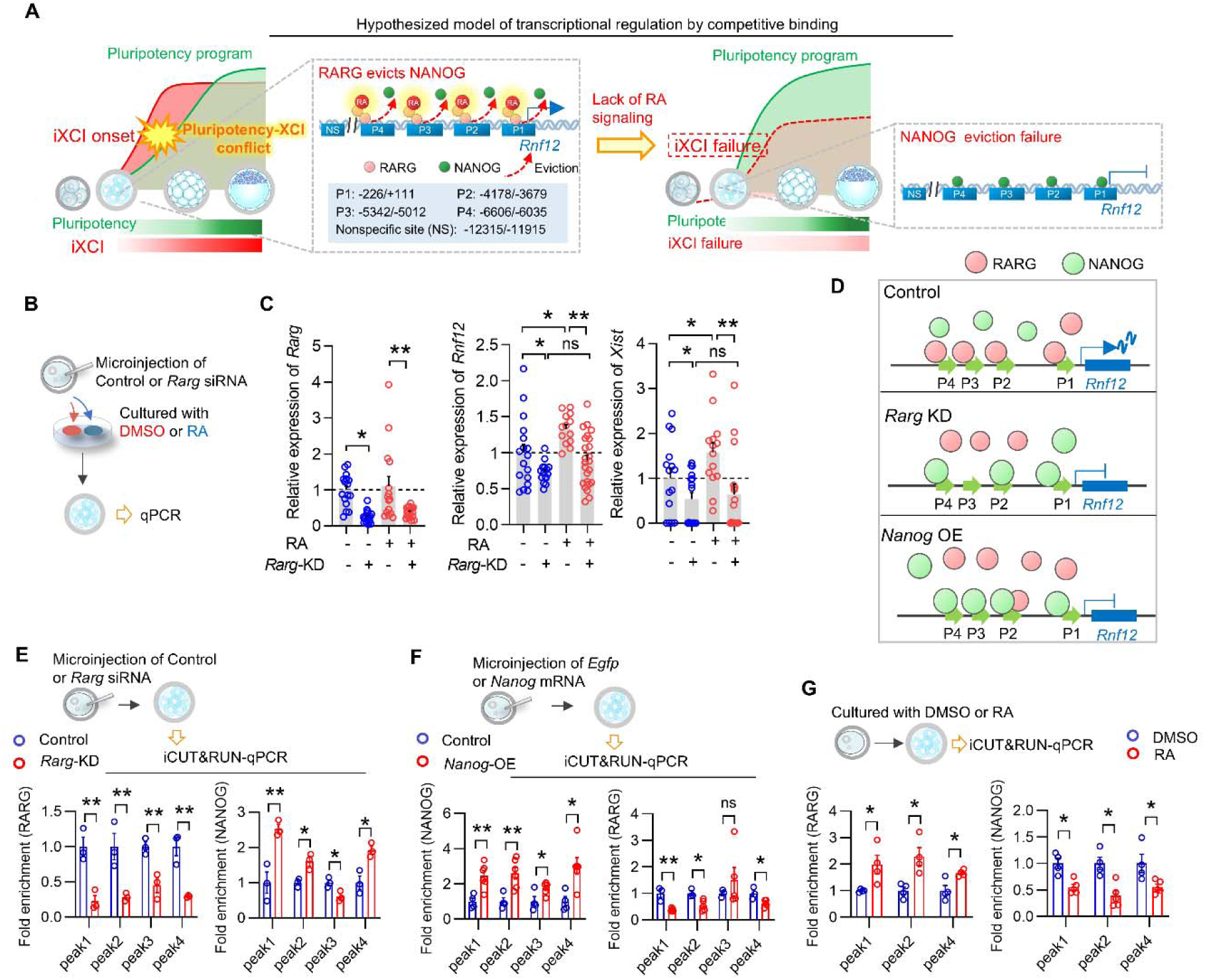
RA activates RARG to evict NANOG from the 5’region of *Rnf12*. **(A)** Schematic diagram of the hypothesis of competitive binding between RARG and NANOG at *Rnf12* 5’ region resolving the conflict between pluripotency acquisition and iXCI onset. **(B)** Schematic diagram of the experimental workflow. *Rarg* knockdown was performed by microinjecting *Rarg* siRNA into zygotes in the presence or absence of exogenous RA. **(C)** Expression levels of *Rarg*, *Rnf12*, *Xist* were detected in female 8-cell embryos subjected to *Rarg* knockdown or RA supplementation, either alone or in combination. Each dot represents a single female 8-cell embryo. **(D)** Schematic diagram of the model in which RARG competes with NANOG for binding to peak1, peak2 and peak4 within *Rnf12* 5’region under the conditions of *Rarg* knockdown and *Nanog* overexpression respectively. **(E)** iCUT&RUN-qPCR analysis of RARG and NANOG occupancy at their putative binding sites within *Rnf12* 5’ region in 8-cell embryos subjected to *Rarg* knockdown or not. **(F)** iCUT&RUN-qPCR analysis of RARG and NANOG occupancy at *Rnf12* 5’ region in 8-cell embryos subjected to *Nanog* overexpression or not. **(G)** iCUT&RUN-qPCR analysis of RARG and NANOG occupancy at *Rnf12* 5’ region in 8-cell embryos exposed to exposed to exogenous RA. In **E-G,** each dot represents a biological replicate (50 8-cell embryos per replicate). Data are mean ± s.e.m. Two-tailed Student’s t-tests. \**p* < 0.05, \*\**p* < 0.01, ns, not significant.

First, to confirm RARG’s role, we deleted *Rarg* using CRISPR/Cas9 in zygotes. *Rarg* knockout significantly reduced both *Rnf12* and *Xist* expression (Figure. S5A), as well as *Xist* RNA coating (Figure. S5B-D). Notably, *Rarg* knockdown also abolished the effect of RA induction (Figure. 3B, C), indicating that RA-induced activation of the *Rnf12*-*Xist* axis is dependent on RARG.

Next, to test the competition between RARG and NANOG for binding to *Rnf12* regulatory elements, we performed *Rarg* knockdown and *Nanog* overexpression respectively and compared their binding enrichment (Figure. 3D). iCUT&RUN-qPCR showed that reducing RARG recruitment via *Rarg* knockdown led to significantly increased NANOG binding to peaks 1, 2 and 4 (Figure. 3E), resulting in *Rnf12* repression (Figure. 3C). Conversely, increasing NANOG recruitment via its overexpression reduced RARG binding to peaks 1, 2 and 4 (Figure. 3F), lowering *Rnf12* expression (Figure. 1C). In line with these, *Rarg* overexpression reduced NANOG binding (Figure. S5E-F) while *Nanog* knockdown increased RARG binding on these elements (Figure. S5G), both of which facilitated *Rnf12* induction (Figure. 2K and Figure. S5H). These findings suggest that RARG activates *Rnf12* by competing with NANOG for binding to the regulatory elements within the *Rnf12* 5’ region.

To explore RA’s influence on this competition, we added exogenous RA to the culture medium. This treatment enhanced RARG binding and reduced NANOG occupancy at peaks 1, 2, and 4 of the *Rnf12* 5’region (Figure. 3G), in line with increased *Rnf12* expression (Figure. 2D), thereby safeguarding *Xist* upregulation.

Together, our results demonstrate that RA stimulates RARG to compete with and evict NANOG from *Rnf12* 5’ regulatory elements, thus preventing NANOG’s repression and ensuring the co-upregulation of *Xist* and *Nanog*. This mechanism resolves the conflict between co-establishment of iXCI and pluripotency program during the early cleavage stage, thus safeguarding iXCI, even in the face of robust *Nanog* upregulation. Our finding not only supports the master coordinator role of RA for orchestrating appropriate progression of both iXCI and pluripotency program, but also highlights the critical role of peaks 1, 2, 4 within *Rnf12*’s regulatory elements in supporting this conflict-resolution mechanism. A key question remains: how essential are these elements for *Rnf12* expression, and how do they achieve their transcription-activating function via RA induction (Figure. 4A)?

**Figure 4.**
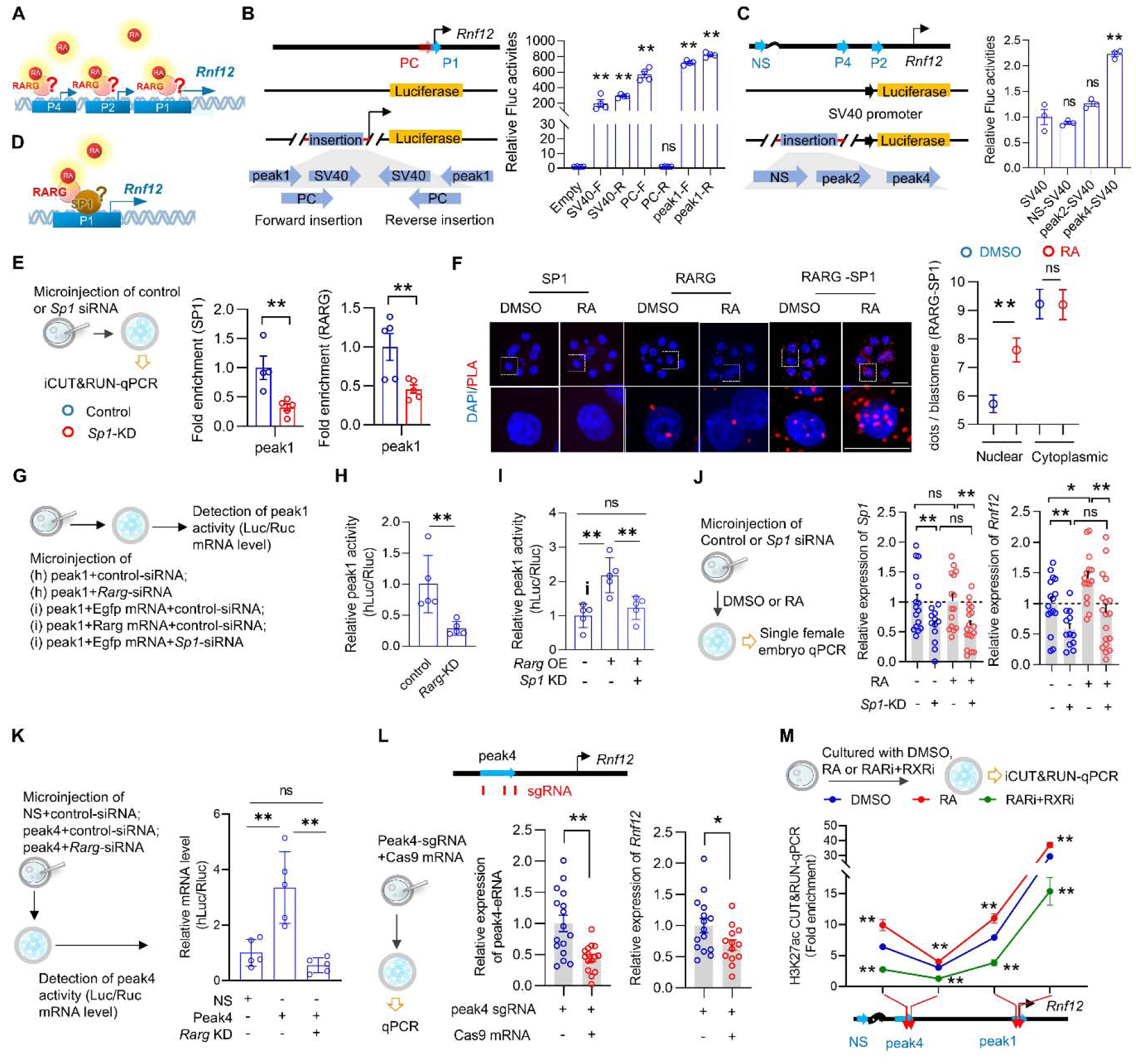
RA induces *Rnf12* dependent on the scaffolding function of SP1 within the 5’region of *Rnf12*. **(A)** Diagram illustrating the unknown mechanisms of RARG binding to peak1, peak2, and peak4. **(B)** Schematic diagram of luciferase reporter constructs containing different forward and reverse-directed promoter segments. Positive control (PC) was the previously identified *Rnf12* promoter ^93^. SV40 is the classical promoter. The promoterless vector was used as the control. **(C)** Schematic diagram of luciferase reporter constructs containing potential enhancer elements. The enhancerless vector was used as the control. NS, non-specific site. Right panels in (B) and (C): relative activities of the presented constructions transfected into HEK293T cells. **(D)** Schematic diagram of the hypothesized model in which SP1 links RARG and stabilizes it at peak1 to induce *Rnf12* expression. **(E)** iCUT&RUN-qPCR analysis of relative SP1 and RARG occupancy at peak1 in 8-cell embryos subjected to *Sp1* knockdown or not. Each dot represents a biological replicate (50 8-cell embryos per replicate). **(F)** Validation of the interaction between RARG and SP1 via in situ PLA in 8-cell embryos exposed to exogenous RA or not. Right panel: quantification of either nuclear and cytoplasmic signals of PLA (red dots) in 8-cell embryos exposed to exogenous RA or not. **(G)** Schematic diagram of the experimental workflow to measure peak1 activity in 8-cell embryos. **(H)** Relative expression levels of firefly luciferase and renilla luciferase (hLuc/Rluc) were measured *Rarg* knockdown, which indicated peak1 activity. **(I)** Relative expression levels of firefly luciferase and renilla luciferase (hLuc/Rluc) were measured *Rarg* overexpression and *Sp1* knockdown. **(J)** Left panel: schematic diagram of the experimental workflow. *Sp1* knockdown was performed by microinjecting *Sp1* siRNA into zygotes in the presence or absence of RA. Right panel: relative expression levels of *Sp1* and *Rnf12* in female 8-cell female embryos subjected to RA supplementation or *Sp1* knockdown, either alone or in combination. Each dot represents a single female 8-cell embryo. **(K)** Schematic diagram of the experimental workflow. Relative expression levels of firefly luciferase and renilla luciferase (hLuc/Rluc) were measured *Rarg* knockdown, which indicated Peak4-enhancer activity in 8-cell embryos. **(L)** Left panel: schematic diagram of the experimental workflow. Cas9 mRNA and three sgRNAs (red bars) were injected into zygotes. Right panel: the effect of CRISPR-Cas9-mediated depletion of peak4 on the expression level of peak4-eRNA (indicates the knockout efficiency) and *Rnf12* in female 8-cell female embryos. Each dot represents a single female 8-cell embryo. **(M)** iCUT&RUN-qPCR analysis of H3K27ac enrichment at peak1 and peak4 in 8-cell female embryos exposed to either exogenous RA or pan-RAR/RXR inhibitors. Data are mean ± s.e.m. Two-tailed Student’s t-tests. \**p* < 0.05, \*\**p* < 0.01, ns, not significant. Scale bar = 25 μm.

### RA-dependent induction of *Rnf12* requires the scaffolding function of SP1 within its 5′ regulatory region

Integrated analyses of open chromatin landscapes in preimplantation embryos and mESCs (Figure. S6A), indicated peak1 as a promoter and peak4 as an active enhancer. These were further validated by luciferase reporter assays (Figure. 4B-C and Figure. S6B). In the canonical retinoid signaling pathway, retinoic acid response elements (RAREs) are typically required for retinoid receptor binding ^18,19^. However, no RARE was found in peak1 (Figure. S6C), suggesting that RARG binding at the *Rnf12* promoter may require an adaptor. To search for the adaptor, we predicted candidate transcription factors that (1) are expressed in preimplantation embryos, (2) have binding motifs within peak1, and (3) potentially interact with RARG (Figure. S6D-E). Among the candidates, we identified *Sp1*, whose critical role in facilitating *Rnf12* expression was supported by knockdown experiments (Figure. S6F).

The interaction between SP1 and RARG was confirmed by coimmunoprecipitation (co-IP) and proximity ligation assay (PLA) in mESCs (Figure. S6G) and 8-cell embryos (Figure. S6H), respectively, suggesting that the RA-RARG-induced activation of peak1 depends on the linking and synergistic action of SP1 (Figure. 4D). To further investigate SP1’s role in recruiting or stabilizing RARG to peak1, we conducted *Sp1* knockdown experiments (Figure. S6I-J), which led to a decrease in RARG binding to peak1 (Figure. 4E). Conversely, *Sp1* overexpression enhanced RARG recruitment (Figure. S6K-L). It should be noted that the observed changes in enrichment were not attributable to alterations in *Rarg* expression itself (Figure. S6J, M). Notably, RA enhanced the SP1-RARG interaction in the nucleus (Figure. 4F), increasing SP1 and RARG binding to peak1 (Figure. S6N). Together, these results indicate that SP1 recruits or stabilizes RARG at peak1 and RA enhances the RARG-SP1 complex formation and its binding to peak1.

Since luciferase reporter assays indicate that either *Rarg* or *Sp1* overexpression could stimulate peak1 activity, with further enhancement by RA signaling (Figure. S6O), we then tested whether RARG and SP1 synergistically regulate the peak1’s transcriptional activity using luciferase reporters directly injected into embryos. Luciferase reporter assays in 8-cell embryos showed that *Rarg* knockdown decreased peak1 activity (Figure. 4G-H), while *Sp1* knockdown diminished the enhancement of RARG on peak1 activity (Figure. 4G and I). In line with this, *Sp1* konckdown blocked RA’s ability to activate *Rnf12*, while *Sp1* overexpression amplified RA’s effect in 8-cell embryos (Figure. 4J and Figure. S6P). These results suggest that RA-RARG-induced *Rnf12* activation is largely dependent on SP1.

Next, we focused on peak4’s role as an enhancer. Luciferase reporter assays in 8-cell embryos indicated that peak4’s transcription-enhancing activity is dependent on *Rarg* (Figure. 4K). In addition, peak4’s transcription-enhancing activity was potentiated by either *Rarg* overexpression or RA exposure (Figure. S6Q). Deleting peak4 via zygotic CRISPR/Cas9 significantly reduced *Rnf12* expression in 8-cell embryos (Figure. 4L). To further determine whether RA signaling regulates peak4 enhancer activity in embryos, we examined enhancer RNA (eRNA) transcription and H3K27ac enrichment levels, which serve as the markers of the active enhancer ^20,21^, at peak4 following RA signaling manipulation. We found that RA signaling promoted H3K27ac enrichment on peak4 and eRNA transcription at peak4, indicating active enhancer function, while RA signaling inhibition decreased peak4-enhancer’s activities (Figure. 4M and Figure. S7A). Within a topological chromatin domain, active enhancers can interact with a promoters through CTCF/cohesin-anchored enhancer-promoter loop (E-P loop) ^22,23^. Hi-C analysis of ESCs and 8-cell embryos revealed that peaks 1 and 4 reside within the same cohesin-mediated topological domain (Figure. S7B-D), suggesting high frequency of enhancer (peak4)-promoter (peak1) interaction depended on RAD21, the core subunit of cohesin ^22^. In addition, re-analyses of Precision Run-On sequencing (PRO-seq) data of mESCs ^24^ revealed that *Rnf12* and peak4 eRNA expression also decreased after the depletion of RAD21, indicating the importance of cohesion-stabilized E-P loop structure for the transcriptional activities of peak4-enhancer (Figure. S7E). Consistent with findings in ESCs, we detected significant enrichment of CTCF and RAD21 at neighboring regions of peak4 in 8-cell embryos (Figure. S7F). Furthermore, disrupting the loop structure through *Rad21* knockdown resulted in decreased *Rnf12* and *Xist* expression (Figure. S7G), as well as reduced *Xist* RNA coating (Figure. S7H-J). Taken together, our results demonstrate that peak4 enhancer and RAD21-mediated E-P loop are essential for RA-induced *Rnf12* transcription. RA’s activator role on *Rnf12* promoter depends on the scaffolding effect of SP1 within a E-P loop (Figure. S7K). As key regulatory elements co-occupied by RARG and NANOG, both peak 1 and 4 are highly sensitive to RA signaling in preimplantation embryos (Figure. 4M).

### Both maternal VA intake and embryonic RA synthesis are indispensable to iXCI and proper female embryogenesis

Having identified the critical role of RA in orchestrating iXCI establishment and pluripotency program using *in vitro* models, we next investigated how RA safeguards iXCI and whether RA differentially affects male and female embryogenesis under physiological conditions. We first profiled the dynamic expression of RA synthetic and degrading enzymes during preimplantation development. We found that *Raldh2* was the major RA synthetic enzyme in embryos (Figure. S8A-C), while *Raldh1* was the major RA synthetic enzyme in oviducts (Figure. S8D). In addition, we detected RA in both embryos and oviduct fluid at each stage during preimplantation development (Figure. S8E-G). Because RA is a highly diffusible small molecule ^18,25^, both maternal RA supply and embryonic RA synthesis should be considered. Thus, we established several combined models to mimic maternal deficiency and embryonic synthesis defects using VA-deficient (VAD) diet and *Raldh2* knockout mice ^26^ (Figure. 1A). We confirmed the successful knockout of *Raldh2* on genomic level and expression level by single embryo PCR and qPCR, respectively (Figure. S8H-I). Further analysis revealed a decrease of RA levels in oviducts and embryos from a VAD model and a VAD *Raldh2*^+/-^ combined model (Figure. S8J), demonstrating the successful establishment of maternal or embryonic RA deficiency models (Figure. S8K).

Quantifications of *Xist* RNA (Figure. 5B, C and Figure. S9A-B) and H3K27me3 domains (Figure. S9C-F) revealed that when wild-type females fed a VAD diet were crossed with *Raldh2*^+/-^ males, maternal RA is deficient (Figure. S8J-K), both wide-type and *Raldh2*^+/-^ female embryos exhibited progressively severe iXCI impairment. *Raldh2*^-/-^ female embryos from *Raldh2*^+/-^ females (VAD diet) crossed with *Raldh2*^+/-^ males showed the most pronounced loss (∼50% cells) of *Xist* RNA and H3K27me3 domains. In contrast, when wild-type females on a normal diet were crossed with *Raldh2*^+/-^ males, the maternal RA supply was normal (Figure. S8J-K), only *Raldh2*^+/-^ female embryos exhibited impaired iXCI which showed defects of *Raldh2* expression in those embryos (Figure. S8I). These findings demonstrate that both maternal RA intake and embryonic RA synthesis, are crucial for the iXCI. Importantly, maternal RA supplementation completely rescued abnormal iXCI, regardless of their genotypes (Figure. 5B-C and Figure. S9).

**Figure 5.**
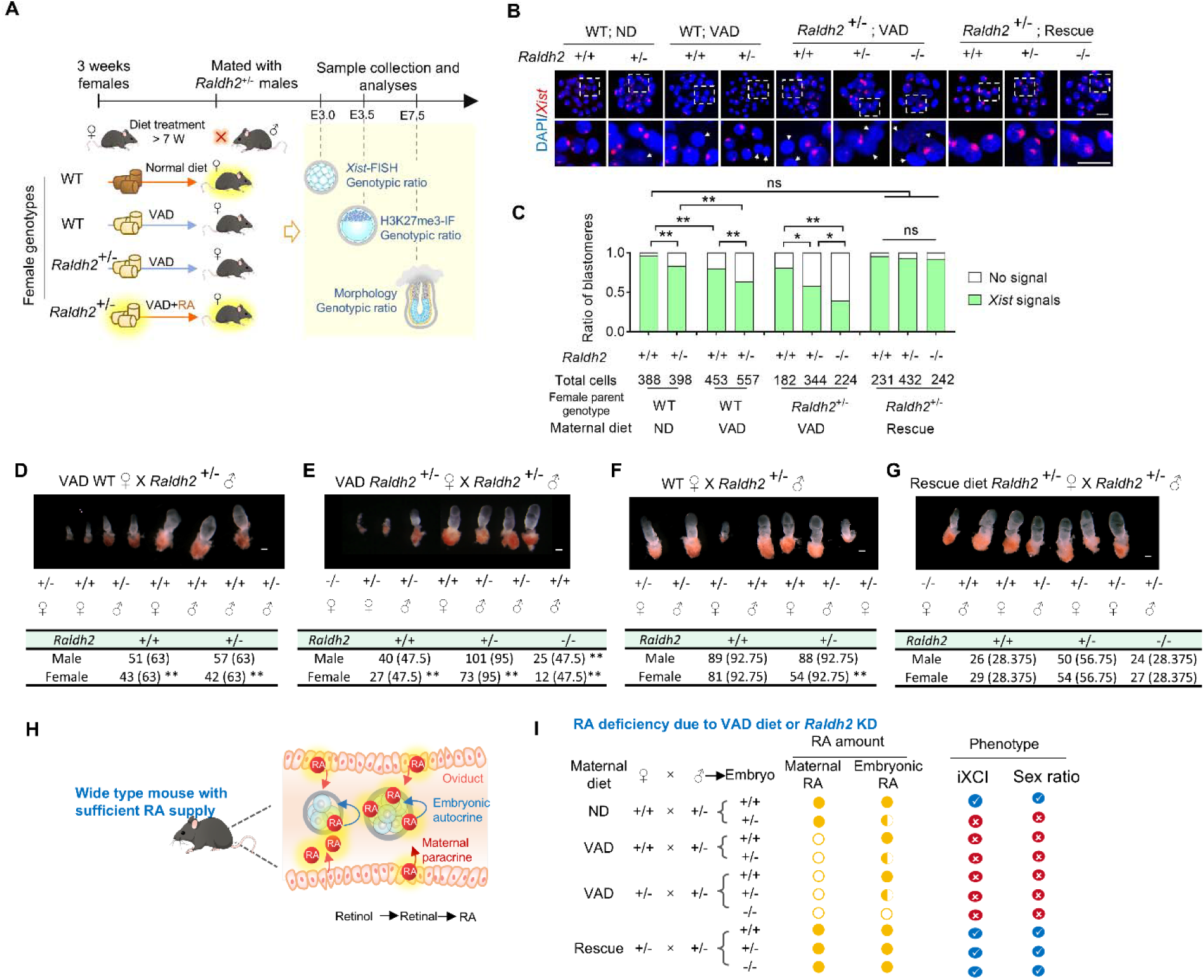
Both maternal VA intake and embryonic RA synthesis are indispensable to iXCI and female embryogenesis. **(A)** Schematic diagram of the experimental design. Combined model recapitulating maternal VA deficiency and embryonic RA synthetic defects was constructed to evaluate the effect of maternal or embryonic RA on female embryos and iXCI status. **(B)** *Xist-*FISH analysis in female morulae from different intersections and maternal dietary conditions. The lowest panel: higher magnification of boxed regions. White arrows indicate blastomeres without the *Xist* RNA domain. Scale bar = 25 μm. **(C)** The percentage of *Xist*-positive and -negative blastomeres to total blastomeres in female morulae. (**D-G**) Representative images (upper panel) and genotype data (lower panel) of embryonic day (E)7.5 female and male embryos from different intersections and maternal dietary conditions. Scale bar = 200 μm. **(H)** Schematic diagram of synergistic supply of RA via both maternal paracrine and embryonic autocrine. **(I)** Summary of phenotypes (iXCI and sex ratio) resulting from RA deficiency caused by parental genotypes or maternal dietary conditions. Chi-square tests were used to determine statistical significance. \**p* < 0.05, \*\**p* < 0.01, ns, not significant.

To rule out secondary effect of compromised oocyte quality due to RA deficiency ^27^, and provide direct evidence of RA’s role in regulating embryonic iXCI, we next used two well-established methods to deplete RA within embryos (Figure. S10A): (1) knockdown of *Raldh2* to inhibit embryonic RA synthesis ^28^ and (2) overexpression of *Cyp26a1*, the main RA-degrading enzyme in preimplantation embryos ^29,30^. When embryonic RA was reduced by *Raldh2* knockdown (Figure. S10B-D), female embryos exhibited lower expression levels of *Xist* (Figure. S10E) and notable loss of H3K27me3 domains (Figure. S10F-G). Likewise, *Cyp26a1* overexpression-induced RA degradation (Figure. S10H-J) also led to significant loss of *Xist* (Figure. S10K-M) and H3K27me3 domains (Figure. S10N-P). Direct supplementation of exogenous RA in embryo culture medium successfully rescued iXCI impairments under both conditions (Figure. S10F-G and K-P).

We next tested the physiological role of maternal VA intake and embryonic RA synthesis on embryogenesis (Figure. S11A). Quantifications of phenotypes showed that crossing wild-type females fed a VAD diet with *Raldh2*^+/-^ males, resulted in a significantly reduced number of both wild-type and *Raldh2*^+/-^ female embryos, deviating from the expected Mendelian ratio (Figure. 5D). *Raldh2*^-/-^ female embryos from *Raldh2*^+/-^ females (VAD diet) crossed with *Raldh2*^+/-^ males had the lowest survival rate, with only about 25% of the expected ratio (Figure. 5E). However, when wild-type females on a normal diet were crossed with *Raldh2*^+/-^ males, only *Raldh2*^+/-^female embryos showed a decrease in survival rate (Figure. 5F). Similar to the rescued iXCI status, the restoration of RA via maternal dietary supplementation fully rescued female embryonic defects and corrected the deviation from the Mendelian ratio caused by either maternal deficiency or embryonic synthetic defects (Figure. 5G, and Figure. S11A). To pinpoint the timing of female developmental defects, we genotyped preimplantation embryos under various genotypic and diet conditions. No deviation from the Mendelian ratio deviation was seen in either morulae or blastocysts (Figure. S11B), suggesting that female developmental defects occur during or shortly after implantation, in line with the developmental defects observed in female embryos with impaired iXCI ^10,31,32^. Finally, both VAD diet model (Figure. S11C-E) and *Raldh2* knockout mice (Figure. S11F-H), resulted in a higher proportion of male offspring.

The prolonged effect of preimplantation RA signaling was also confirmed by transferring the nuclear receptor signaling-inhibited blastocysts to normal recipients for further development (Figure. S12A-C), as well as via intraperitoneal injection of RAR/RXR inhibitors during the preimplantation window (Figure. S12D-N), both of which resulted in female-biased developmental defects.

These data provide *in vivo* evidence that iXCI and female embryogenesis are highly sensitive to physiological RA levels, which are synergistically supported via both maternal paracrine and embryonic autocrine (Figure.5H). Insufficient RA supply due to the maternal intake deficiency or impaired embryonic synthesis, leads to impaired iXCI and female-biased developmental defects, therefore skewing offspring sex ratio at birth (Figure. 5I).

Taken together, our findings support a complex multi-layer model in which VA-RA signaling orchestrating iXCI establishment and pluripotency program during the in cleavage stage of female preimplantation embryos, to resolve the encounter of two conflicting events: RARG not only directly activates the *Rnf12* expression, but also synergizes with RXRA to co-suppress excessive *Nanog*, thus derepressing the *Rnf12*-*Xist* axis; More directly, RA activates RARG to evict NANOG from *Rnf12* 5’ regulatory elements, serving as a conflict-resolution mechanism (Figure. 6).

**Figure 6.**
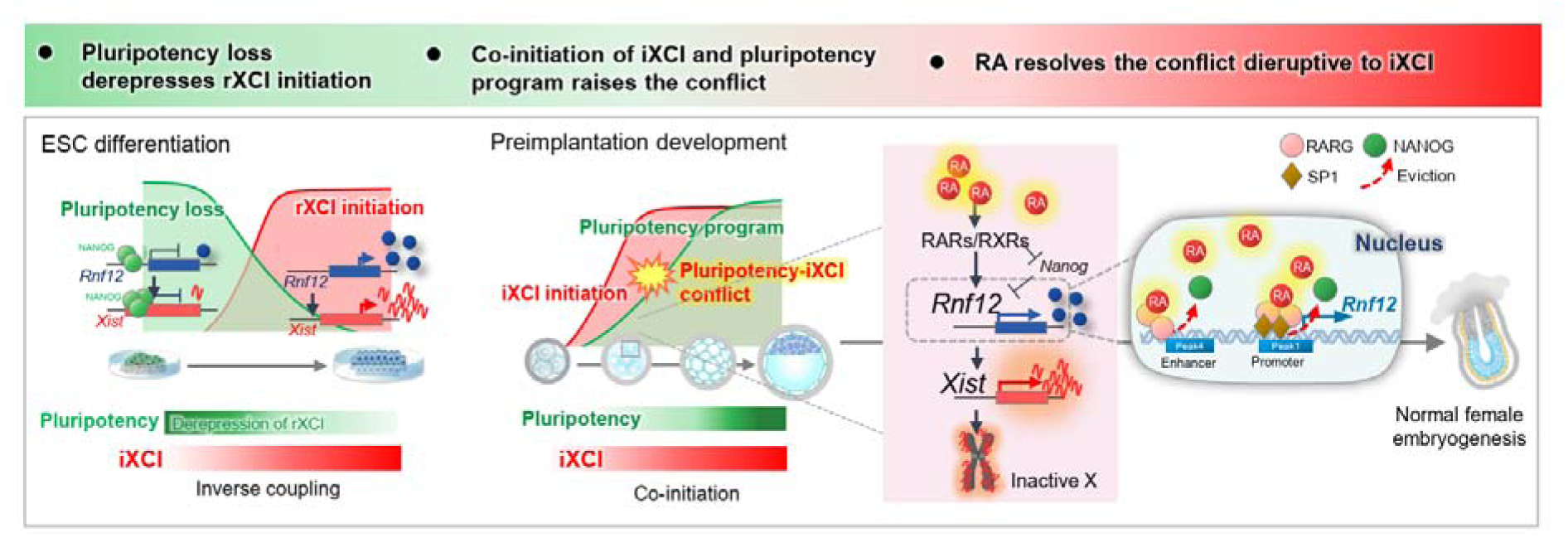
Model illustrating the role and mechanism of RA in resolving the conflict between iXCI and pluripotency program in female cleavage-stage embryos.

## DISCUSSION

XCI and pluripotency are tightly coupled and their spatiotemporal coordination during the differentiation of mouse ES cells and epiblasts, is a classical paradigm of holistic developmental regulation. However, the co-establishment of pluripotency program and iXCI during early cleavage stages challenges the holistic pattern, but how the two are interconnected has been long considered controversial. Although it has been proposed that NANOG and other pluripotency factors do not repress iXCI until the inactive X-chromosome is reactivated in ICM, termed X-chromosome reactivation (XCR) ^6,33,34^, our data definitely indicate that NANOG is the major repressor of iXCI establishment. Given NANOG’s key role in pluripotency acquisition ^9^, RA-induced repression of *Nanog* is unlikely to be the main mechanism to resolve the conflict. Instead, our study reveals that the antagonistic interaction between RARG and NANOG at the *Rnf12* 5’region, along with RA-induced eviction of NANOG, ensures the co-upregulation of *Nanog* and *Xist*.

VA is a vital nutrient and VA deficiency is among the most prevalent nutrition-related health issues globally ^35,36^. As a critical morphogen in chordates ^18^, VA and RA are known to regulate many fundamental biological processes ^18,37–39^. Although RARG binding motifs and expression are strongly enriched during the 2-cell to 8-cell stages ^40^, the role of RA signaling in preimplantation embryos has remained largely unclear, despite recently reported prompting effect on zygotic genome activation during the totipotency window ^15^. Our study fills this gap by identifying a novel role of VA-RA signaling that underlies a critical conflict-resolution mechanism, thus safeguarding iXCI establishment in cleavage-stage embryos. While previous studies have identified many intrinsic regulators of XCI, such as transcriptional activators and noncoding RNAs ^6,7,16,41–43^, extrinsic factors like nutrients have been underexplored. Our results expand the mechanistic understanding of XCI regulation, particularly in relation to nutrient availability.

Our study also highlights the key role of SP1, a general transcription factor, in regulating iXCI. SP1 is traditionally associated with basal transcription of genes involved in essential cellular processes like nucleic acid metabolism and cell cycle regulation. However, recent evidence suggests that SP1 forms transactivation complexes with other transcription factors, fine-tuning gene expression in response to physiological and pathological stimuli, such as in carcinogenesis and neurogenesis via interactions with estrogen and nuclear retinoid receptors ^44–46^. We propose that SP1 may mediate the basal transcription of the *Rnf12*-*Xist* axis, while RARG enhances its transcription through direct interaction and synergy with SP1 during the iXCI establishment window in female embryos. This may explain why some female cells can still establish iXCI even in the absence of RA or receptor function.

Our findings highlight RA as a master regulator ensuring proper female embryogenesis. By elucidating the conflict-resolution mechanism, we also established a mechanistic link between a vital nutrient and offspring sex ratio, explaining the intriguing observation that regions with high prevalence of VA deficiency show a higher proportion of male births ^35,47^. The sex ratio at birth in humans and mammals is an important demographic and ecological indicator, often skewed by many influential factors, such as maternal nutrition ^48–51^, paternal age ^52–54^, male fertility ^55^, and exposure to environmental toxins or stress ^56,57^. However, up to now, the mechanisms of skewed sex ratio in mammals remain unknown. Our findings also provide a framework for understanding other nutritional or environmental factors that may influence sex ratio by affecting sex-specific epigenetic regulation and development.

While a previous report has shown that *Raldh2*^-/-^ embryos are smaller and have severe developmental defects, such as axial rotation disruption and disorganized extraembryonic vessels at embryonic day (E)9-11, leading to death at mid-gestation ^58^. Our data reveal a previously unrecognized sex dimorphic pattern of RA deficiency-induced developmental defects: either *Raldh2*^-/-^ or *Raldh2*^+/-^ female embryos showed earlier and more severe defects or lethality than their male counterparts. Our finding aligns with the reports that cord plasma VA levels are higher in female neonates compared to males ^59,60^, suggesting that female embryos require a higher threshold of RA for development. The sexually dimorphic effects of RA are supported by increasing evidence in metabolic regulation ^61^, gametogenesis ^62^ and other morphogenetic processes ^63,64^. RA is a well-studied molecule and a potent clinical drug, used in treatments for various cancers, including acute promyelocytic leukemia, ovarian cancer, and breast cancer ^65^. XCI also plays a critical role in several diseases, including autoimmune diseases ^66^, ovarian cancer ^67–69^, and breast cancer ^70–72^. The newly discovered link between RA signaling and XCI in our study opens the possibility for developing diagnostic and therapeutic interventions targeting VA deficiency and XCI-related diseases. More broadly, our findings underscore the sexually dimorphic effects of maternal nutrition, with profound implications for gender-specific clinical applications.

### Limitations of the study

In this study, we reported that RA ensures proper female embryogenesis via safeguarding iXCI, and resolves the conflict between pluripotency acquisition and iXCI onset via competitive binding between RARG and NANOG at Rnf12 5’ region. However, several key questions warrant further investigation. First, although we showed that Xist expression, Xist RNA coating, and H3K27me3 domains were all impaired by RA signaling blockage, we have yet to profile how many X-linked genes would be affected. Also, it would be more informative to elucidate whether RA receptors compete with NANOG on a genome-wide scale in both male and female embryos to orchestrate proper embryogenesis. Although we showed that RA is required for successful iXCI establishment, whether RA is required for rXCI and how RA affects rXCI need further study.

## STAR Methods

### Animal Studies

All ICR and C57 mice used in this study were obtained from SPF Biotechnology Co. Ltd. Heterozygous *Raldh2*^+/-^ mice were obtained from the Mutant mouse research and resource center (ID 41427, MMRRC) and maintained in a C57BL/6 background ^26^. Genotypes and sexes were determined by PCR. Primers are listed in **Table S1**. All mice were maintained in a climate-controlled room on a 12 hours light/dark cycle and allowed food and water ad libitum. All animal experiments were approved by the China Agricultural University Institutional Animal Care and Use Committee and performed in accordance with the committee’s guidelines.

### Vitamin A free diet and RA supplementary diet

To construct maternal vitamin A deficiency mice, 3-weeks old C57 female mice were kept in a AIN 93G vitamin A-free diet (Huafukang biological technology Co., Ltd, Beijing, China) ^73^ at least 7 weeks as previously described ^74–77^. To construct RA rescue models, 3-weeks old female mice were kept in a rescue diet, which was the VAD supplemented with RA 0.1mg/g food ^26,78–81^. Embryos were collected from pregnant female mice under normal diet, VAD diet, or rescue diet for further analysis.

### Preparation and collection of mouse embryos

All experiments involved in embryo preparation were performed as previously described ^82^, with minor modifications. Female mice (7-8 weeks) were super-ovulated through an initial injection of 5 IU pregnant mare serum gonadotropin (PMSG, Ningbo Hormone Product Co., Ltd, Ningbo, China), followed by a 5 IU injection of human chorionic gonadotropin (hCG, Ningbo Hormone Product Co., Ltd) 48 hours later. Super-ovulated female mice were mated with male mice of 10-week-old, and successfully mating was confirmed by the presence of vaginal plug on the next morning. For collecting in vitro cultured (IVC) embryos. Zygotes were collected from the ampullae at 18 h post hCG injection, then cultured in potassium simplex optimization medium containing amino acids (KSOM+AA; Millipore) with or without chemical treatment (DMSO, RA (Sigma, R2625), RAR inhibitor (BMS493, Tocris, 3509) plus RXR inhibitor (UVI3003, Tocris, 3303), MEK inhibitor (PD0325901, Selleck, S1036)) under mineral oil at 37 °C in 5% CO2. 4-cell-, 8-cell-, morula-, blastocyst-stage embryos were collected based on their morphology and developmental time. For collecting in vivo (IVO) embryos, 8-cell-, morula-, blastocyst-stage embryos were collected at 62-66 h, 74-78 h, 96h post hCG injection, respectively. For embryo transfer, blastocysts were transferred to the uteri of Day 3.5 pseudopregnant females. For E7.5 embryo collection, the conceptuses covered with the decidual mass were gently teased away from the uterus. The decidua in which the conceptus embedded was peeled off, and the parietal yolk sac was opened to expose the visceral yolk sac endoderm layer ^10,83^.

### In vitro transcription and microinjection

For mRNA samples, *Cyp26a1*, *Rnf12*, *Nanog*, *Rarg*, *Sp1*, and enhanced green fluorescent protein (*Egfp*) were synthesized by a T7 mMESSAGE mMACHINE Kit (Invitrogen, AM1345) using primers in **Table S1** and then mRNAs were recovered by MEGAclear™ Kit (Invitrogen, AM1098) and dissolved in nuclease-free water to a final concentration of 200 μg/mL. For sgRNA samples, Rarg and Rnf12_peak4 sgRNA were synthesized by a MEGAshortscript™ Kit (Invitrogen, AM1354). For knockdown of *Raldh2*, *Nanog*, *Rarg*, *Klf4*, *Sp1*, and *Rad21*, small interfering RNAs were injected into zygote followed by in vitro culture in KSOM with or without treatment (DMSO, RA, and RARi + RXR). siRNA and sgRNA targeted sequences are included in **Table S1**. Microinjection of mRNA, sgRNA, or siRNA was performed as previously described ^10,83^. Samples were injected at 5–10 pl per zygote. For the knockdown experiment, 100 μM siRNA was used for each siRNA and non-targeting siRNA as a control. 50 ng/μL of each sgRNA and 100 ng/μL Cas9 mRNA were used for knockout experiment.

### Cell culture and differentiation

Undifferentiated PGK12.1 female mES cells were cultured on 0.1% gelatin coated plates with LIF condition ^84^. PGK12.1 mES Cells were grown in stem cell medium (Knockout DMEM base medium (Gibco, 10829018), supplemented with 10% fetal bovine serum (FBS) (Hyclone, SH3007003), 2 mM GlutaMAX (Gibco, 35050061), 1x EmbryoMax® MEM, Non-Essential Amino Acids (Millipore, TMS-001-C), 0.084 mM 2-mercaptoethanol (Gibco, 21985023), 100 IU/ml penicillin-streptomycin (Gibco, 15140122), 1000 IU/ml LIF (Millipore, ESG1107)). To induce differentiation, 3.0∼5.0 × 10^3^/cm^2^ cells were plated onto glasses coated by 5 μg/cm^2^ fibronectin in stem cell medium. After 24 h growth, cells were rinsed by 1 ml DPBS and differentiation medium (N2B27 medium with 5 mg/mL BSA) was added into each well to induce cell differentiation. Cells were incubated in a humidifier containing 5% CO_2_ at 37 °C.

### Quantitative real-time PCR (qRT-PCR) analysis

qRT-PCR of single embryo was performed as previously described ^10^. For single 4-cell or 8-cell embryo, the embryo was collected and treated with 0.5% pronase (sigma, P5147) to remove zona pellucida, then one blastomere was collected in 2 μL ice-cold cell lysis buffer (Thermo Fisher, AM872) for subsequent sex determination, other blastomeres were collected in 5 μL ice-cold cell lysis buffer for subsequent qRT-PCR. For single morula embryo, embryo was collected and treated with 7 μL ice-cold cell lysis buffer. Next, 2 μL lysis product was used for subsequent sex determination, and the remaining 5 μL lysis product was used for subsequent qRT-PCR. For sex determination, 2 μL lysis product was digested with 2.5 μL 1 mg/mL proteinase K (Merck, 1245680100) at 55 °C, 2 h; 85 °C, 15 min. Then, 4.5 μL product was used for PCR with TaKaRa Ex Taq Hot Start Version (TaKaRa, RR006A) and sex determination primers **(Table S1)**. 5 μL cRNA in the lysis product was reverse-transcribed into complementary DNA (cDNA) with a HiScript II reverse transcriptase reagent kit (Vazyme, R223-01, Nanjing, China). Real-time PCR was performed using SsoFast EvaGreen Supermix (Bio-Rad, 1725201) in a CFX96 real-time PCR machine (Bio-Rad). Gapdh and H2afz were used as reference. The primers used for qRT-PCR are listed in Supplementary **Table S1**.

### Immunoprecipitation and western blot

Co-immunoprecipitation (Co-IP) was performed as previously described ^85^ by using Dynabeads™ Protein G Immunoprecipitation Kit (Invitrogen, 10007D, USA). Total cell lysates for each sample were collected from differentiated PGK12.1. anti-SP1 (ABCAM, ab227383) was used for immunoprecipitation. For immunoblot experiments of mouse embryo, 100 mouse embryos per stage were denatured from Laemmli sample loading buffer (Bio-Rad Laboratories) with 5% β-mercaptoethanol (Sigma Aldrich) and protease inhibitor at 100°C for 10 min and stored at −80°C for future use. Western blotting was performed as described previously ^10^. Primary antibodies against GAPDH (1: 2000, Cell Signaling Technology, 5174S), RALDH2 (1:200, Santa Cruz Biotechnology, sc-393204), SP1 (1:200, Santa Cruz Biotechnology, sc-17824 X), RARG (1:200, Santa Cruz Biotechnology, sc-7387 X) were used in this study. The blots were visualized using the ECL system (Bio-Rad, Hercules, CA, USA). The grey level of each band was calculated by ImageJ (https://imagej.nih.gov/ij/, NIH, Bethesda, MD, USA).

### Immunofluorescence

Embryos were fixed in 4% paraformaldehyde at 4 overnight and permeabilized in 0.5% Triton X-100 - 0.1% Polyvinylalcohol - PBS (PBST-PVA) at room temperature for 1 h. After blocking with 1% BSA in PBST-PVA, the samples were incubated in primary antibodies diluted in PBST-PVA containing 1% BSA overnight at 4 . Antibodies were used as follows: anti-RALDH2 (1:200, Santa Cruz Biotechnology, sc-393204), anti-H3K27me3 (1:1000, Millipore, 07-449), anti-CDX2 (1:500, BioGenex, MU392A-UC), anti-RNF12 (1:200, abnova, H00051132-M01), anti-NANOG (1:1000, ABCAM, ab80892), anti-RARG (1:400, Santa Cruz Biotechnology, sc-7387 X), anti-RARG (1:400, ABCAM, ab97569), anti-SP1 (1:400, ABCAM, ab227383), anti-SP1 (1:400, Santa Cruz Biotechnology, sc-17824 X). Following three washes with PBST-PVA, the embryos were incubated with appropriate Alexa 488 or 594 fluorophores conjugated secondary antibodies (A-11012 or A-11029, Thermo Fisher Scientific) for 1 h at room temperature. Finally, the samples were counterstained with DAPI. The fluorescence signals were imaged using an BX51 microscope (Olympus). The fluorescence signals were imaged using a confocal laser scanning microscope (Digital Eclipse C1; Nikon). The fluorescent intensity was calculated using ImageJ.

### ELISA for RA detection

RA concentrations using an ELISA colorimetric detection kit according to the manufacturer’s instructions (MyBioSource, MBS705877). For embryos, 50 embryos of each replicate were used for RA concentration detection. The detection of RA content in oviductal fluid at each stage was performed according to the published protocol ^86^. Then, the concentration was calculated by dividing by the relative volume of oviductal fluid at each stage ^87,88^. This kit could not distinguish all-trans retinol, all-trans retinal or all-trans retinoic acid. Thus, the concentration of RA was the concentration of all forms.

### LC-MS/MS for RA

For the detection of RA in uterus and oviduct, all-trans retinoic acid was extracted from snap frozen mouse uterus and oviducts as described ^89^. Quantification of RA was performed by the MetWare (Wuhan, China) using the SCIEX QTRAP 6500+ liquid chromatography tandem mass spectrometry / tandem mass spectrometry (LC-MS/MS) system.

### RNA FISH

The ViewRNA ISH Cell Assay kit (Invitrogen, AVC0001) was used for RNA-FISH, and the procedure was performed as previously described ^10,90,91^. Fixed embryos were treated with Detergent Solution QC at room temperature for 5 min, and digested with protease QS at room temperature for 10 min. Next, the embryos were hybridized with probes against *Xist* or *Rnf12* (Invitrogen). Then the embryos were hybridized with PreAmplifier and Amplifier Mix, respectively. Label Probe Mix was used to produce a signal. The embryos were counterstained with DAPI, and imaged using an attached digital microscope camera (DP72; Olympus).

### Reporter assay in HEK293 and embryos

For RARE-SV40-DsRed plasmid construction, DsRed-monomer fragment from pDsRed-Monomer-N1 plasmid was cloned to replace luciferase fragment in pGL3-RARE-luciferase ^92^ (13458, Addgene) to reconstruct RARE-SV40-DsRed reporter vector, whose sensitivity was verified in HEK293 (Figure. S2A). Then RARE-SV40-DsRed and SV40-Egfp fragments was amplified and purified for pronuclear microinjection.

PC (-667/+1), Peak1 (-226/+111), NS (-12315/-11915), SV40, peak2 (-4178/-3679) -SV40, and peak4 (-6606/-6035)-SV40 were cloned into the luciferase reporter pEZX-FR01. Positive control (PC) was the reported mouse *Rnf12* promoter ^93^. For the reporter assay in HEK293, the inserted luciferase reporter and pCAGGS-Rarg or pCAGGS-Sp1 overexpression plasmid were transfected to HEK293 with lipofectamine 2000 (Invitrogen, 11668019). Luciferase activity was measured at 24 h after transfection using the Dual-Luciferase Reporter Assay System (Promega) following the manufacturer’s protocol. In regard to plasmids used for *Rarg* or *Sp1* overexpression in HEK293, Rarg or Sp1 complementary DNA were cloned into pCAGGS vector. For the detection of peak1 and peak4 activity in 8-cell embryos, the fragments of peak1-firefly luciferase, peak4-SV40-firefly luciferase, NS-SV40-firefly luciferase, and SV40-renilla luciferase were amplified and purified for microinjection (20 ng/μl) into zygotes. 8-cell embryos were collected for qRT-PCR of firefly luciferase and renilla luciferase mRNA level. Renilla luciferase was used as reference.

### Proximity ligation assays

The in situ proximity ligation assays (PLA) was performed using the Duolink® In Situ Red Starter Kit according to the manufacturer’s instruction (Sigma, DUO92101). Embryos were fixed and permeabilized as described for immunofluorescence. After blocking, embryos were incubated with primary antibody. Embryos were then incubated with PLA probes, followed by ligation and amplification reaction. Finally, embryos were counterstained with Duolink® In Situ Mounting Medium with DAPI (MilliporeSigma). Furthermore, SP1 (ABCAM, ab227383) and RARG (Santa Cruz Biotechnology, sc-7387 X) single primary antibody alone were used as technical negative controls.

### Single embryo RNA-seq library preparation and sequencing

Single 8-cell embryo was collected and treated with 0.5% pronase (sigma, P5147) to remove zona pellucida, then one blastomere was collected for sex determination as described above, other blastomeres were collected and stored at -80 for later processing. The female 8-cell embryos were used for RNA-seq library construction as previously described ^91,94^. In short, single female embryo was lysed and mRNAs were reverse-transcribed. After amplification, cDNA 3’ terminal was labeled by biotin and then fragmented. cDNA library was constructed by KAPA Hyper Prep Kit (KK8504, KAPA Biosystems, USA). Sequencing was performed on Illumina HiSeq 4000.

### iCUT&RUN-qPCR and data analyses

iCUT&RUN-qPCR assays were performed as previously described with minor modifications ^91,95^. In short, 8-cell embryo samples (total volume with buffer less than 1 μl) were prepared freshly into a 1.5 ml low-binding tube. The zona pellucida was removed with Tyrode’s solution and the polar body removed with a sharp glass pipette. 300 μL cold nuclear extraction buffer (20 mM pH 7.9 HEPES-KOH, 10 mM KCl, 0.5 mM Spermidine, 0.1%Triton X-100, 20% Glycerol, freshly added protease inhibitors). 10 μL Concanavalin A beads (86057, Polysciences, Warrington, USA) were prepared per sample. Beads were washed and resuspended in 150 μL binding buffer (20 mM HEPES-KOH, pH 7.9, 10 mM KCl, 1 mM CaCl_2_, 1 mM MnCl_2_), then mixed with nuclei and incubated for 10 min at room temperature. Bead-bound nuclei were blocked with 1 mL cold blocking buffer (20 mM HEPES, pH 7.5, 150 mM NaCl, 0.5 mM Spermidine, 0.1% BSA, 2 mM EDTA, 0.5 mM Spermidine, freshly added protease inhibitors) for 5min at room temperature. Then samples were placed on a magnetic stand and resuspended in 250 μL cold wash buffer (20 mM HEPES, pH 7.5, 150 mM NaCl, 0.5 mM Spermidine, 0.1% BSA, freshly added protease inhibitors). Primary antibody RARG (Santa Cruz Biotechnology, sc-7387 X), NANOG (Active Motif, 39843), SP1 (ABCAM, ab227383), RAD21 (ABCAM, ab992), CTCF (ABCAM, ab188408), and H3K27ac (Cell signaling technology, 8173) antibody was added to a final concentration of 1:100 at 4 for 2h. IgG (3493998, Invitrogen) was used as negative control. Protein A-micrococcal nuclease (pA-MNase; 0.95 μg/μL pA-MNase was a gift from Wei Xie’s lab (Tsinghua University, China)) was added to a final concentration of 1:750 at 4 ◦C for 1h. Samples were resuspended in 150 μL cold wash buffer and added 3 μL 100 mM CalC2 for 30min. The reaction was stopped with 150 μL 2 × stop buffer (200 mM NaCl, 20 mM EDTA, 4 mM EGTA, 50 μg/mL RNaseA, 40 μg/mL glycogen) at 37 ◦C for 20min, and 3 μL 10% SDS (L4509, Sigma-Aldrich, City of Saint Louis, USA) and 2.5 μL 20 mg/mL Proteinase K (DE102-01-AA, Vazyme) were added to the supernatants incubated at 50 for 1h. DNA was purified using phenol/chloroform/isoamyl alcohol (1328-K169, Amresco, New York, USA) and then used for RealTime-qPCR analysis. The primers are shown in **Table S1**.

For data analyses, we performed as previously described with minor modifications ^91,96^. Briefly, IgG was first used as negative control in Figure. 2B and 2G, which also suggests RARG, NANOG, and RXRA antibodies works well. To avoid the batch effects, the non-specific site (NS), which showed no binding of RARG and NANOG, was used as negative control in the subsequent analyses.

### Data analyses

#### RNA-seq data processing

Methods for RNA-seq read alignments and differential gene expression analysis were similar to previously described ^83^. RNA-seq reads were trimmed and then mapped to the mm10 genome by STAR 2.7.3a ^97^. FeatureCounts v.2.0.1 ^98^ was used to count mapped reads. The genes expressed (count > 0) in more than three samples were retained. Differentially expressed genes (DEGs) were identified with adjusted P < 0.05 and FC > 1.5 by DESeq2 v.1.24.0 ^99^. The gene expression level was normalized to transcripts per million (TPM) (**Table S3**).

#### ChIP-seq data and ATAC-seq data processing

ChIP-seq and ATAC-seq data were mapped to mm10 genome by Bowtie2 (v 2.4.1) ^100^. Duplicate reads were filtered out using picard MarkDuplicates (v2.18.27) ^101^. Genome coverage bigwig files for IGV browser were generated by deeptools (v 3.4.3) ^102^.

#### Sequence Analysis of DNA Binding Motifs

FIMO function (https://meme-suite.org/meme/tools/fimo) ^103^ was used to screen the potential transcription factors (p value<0.001, score>1) on Rnf12 promoter regions (peak1). JASPAR ^104^ was used to screen the potential RARG binding sites on peak1, peak2, and peak4.

### Data, materials, and code availability

All data are available in the main text or the supplementary materials. All data are available within the article and its Supplementary Tables. Accession codes of the published data in GEO used in this study are as follows: ATAC-seq of 8-cell embryos and mES cells, GSE66581 ^40^; RNA polymerase II stacc–seq of 8-cell embryos and mES cells, GSE135457 ^105^; 8-cell H3K4me3, GSE73952 ^106^; 8-cell H3K27ac, GSE72784 ^107^; NANOG ChIP-seq of mES cells, GSE44764 ^108^; RAR and RXR ChIP-seq of RA treated 48h cells, GSE56893 ^109^; RNA-seq data of female mES cells during X-chromosome inactivation, GSE116649 ^110^; RNA-seq data of female pre-implantation embryos, GSE71442 ^4^; PRO-seq, HiC, and RAD21 ChIP-seq of RAD21-depletion and control mES cells, GSE144834 ^24^; H3K4me1, H3K4me3, and H3K27ac of mES cells, GSE80049 ^111^; mES cells p300 ChIP-seq, GSM2417169 ^112^. HiC of 8-cell embryos, GSE82185 ^113^; CTCF ChIP-seq data of mES cells, GSE144056 ^114^. RNA-seq data of bovine embryos, GSE52415 ^115^. RNA-seq data of human embryos, E-MTAB-3929 ^116^. Software used to analyze these data are listed in Methods and are all publicly available.

## Acknowledgments

We thank members of the Jianhui Tian’s lab for discussion and comments during this Retinoic Acid study and preparation of the manuscript. We thank Fuchou Tang and Shuai Gao for providing help in single embryo RNA-seq experiments. We thank Wei Xie, Jing Ma, and Lijia Li for generously providing help in improving our ATAC-seq and CUT&RUN methods. We thank Sarah J. Hainer for generously providing help in improving our CUT&RUN method. We thank Neil Brockdorff and Ingolf Bach for generously providing PGK12.1 female ES cell line. We thank Zhen You for suggestions in bioinformatic analysis. We thank Wei Xie, Zhiyuan Chen, Zhenhai Du, Rocío Melissa Rivera, Chao Wang, Teng Zhang, Hua Zhang, and Yulei Wei for reading the manuscript and providing helpful comments.

## Author contributions

Conceptualization: J.T., L.A. and Q.Y. Experimental designs: Q.Y., L.A., J.T., C.Z., L.M., Y.Y., S.Z. Raldh2 knockout mice breeding and maintenance: C.Z., S.Z., and W.W. Vitamin-A deficiency experiments: H.D. and S.Z. Microinjection of embryos: X.W., L.M., and S.Z. Embryo transfer: Z.Z. Performed RA receptor inhibition experiments and Cyp26a1 overexpression experiments: C.Z. Performed Rarg knockout experiments, Sp1 knockdown experiments, and Rad21 knockdown experiments: L.M. Single embryo RNA-seq library preparation: Q.L. Performed luciferase reporter assay: S.Z. and W.W. Performed iCUT&RUN-qPCR experiments: Q.Y., L.M., and Y.Y. Performed CUT&Tag and RNA-seq of single embryo from Vitamin-A deficiency and Raldh2-knockout mice: S.Z. Performed PLA assay: Y.Y. Performed Co-IP assay: X.H. Performed post-implantation embryo dissection and sex determination: Q.Y., Z.Z., C.Z., and J.L. Performed other embryo experiments: Q.Y., W.W., S.Z., Y.Y., C.Z., J.L., M. Q., Y.T., and W.F. Data interpretation and analysis: Q.Y., S.Z., C.Z., Y.Y, L.A., and J.T. Figures organization: Q.Y., S.Z., and L.A. Writing, reviewing and manuscript editing: L.A., Q.Y., S.Z., and J.T. Revising manuscript: J.T., and J.W. Funding: J.T. and L.A.

## Competing interests

Authors declare that they have no competing interests.

Supplementary Materials

Figs. S1 to S12

Tables S1 to S3

