## Supplementary material for "Retinoic acid resolves the conflict between X-chromosome inactivation and pluripotency program in female cleavage-stage embryos": Figs. S1 to S12

### Supplementary Materials

**The PDF file includes:**

Figs. S1 to S12
References

**Other Supplementary Materials for this manuscript include the following:**

Table S1 to S3

**Fig. S1.**

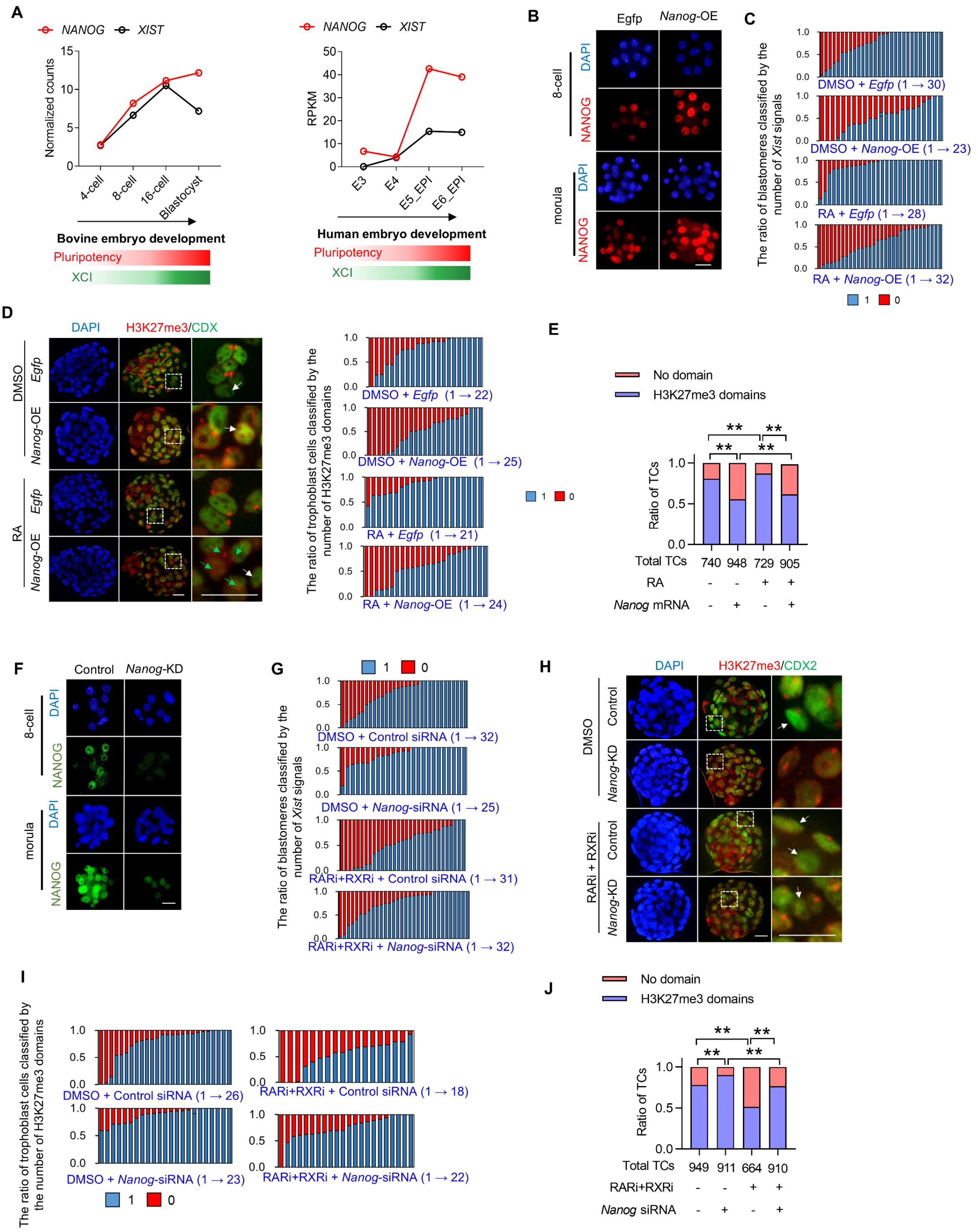


**Figure S1. RA protects iXCI from NANOG's repression. (A)** Expression dynamics of *RNF12*, *XIST* and *NANOG* during bovine (*1*) (left panel) or human (*2*) (right panel) preimplantation development. **(B)** Immunostaining of NANOG in female 8-cell embryos and morulae after overexpression of *Nanog*. **(C)** The ratio of blastomeres classified by the number of *Xist* domains in female morulae subjected to RA supplementation or *Nanog* overexpression, either alone or in combination. Each bar represents one female morula. **(D)** Immunostaining for H3K27me3 (red) in the nuclei (DAPI) of female blastocysts colabeled with CDX2 (green)-positive trophoblast cells. Rightmost panels: higher magnification of boxed regions. White arrows indicate H3K27me3 domain-negative trophoblast cells. Right panel: the ratio of trophoblast cells classified by the number of H3K27me3 domains. Each bar represents one female blastocyst. **(E)** The percentage of H3K27me3-positive and -negative trophoblast cells to total trophoblast cells in female blastocysts. **(F)** Immunostaining of NANOG in female 8-cell embryos and morulae after knockdown of *Nanog*. **(G)** The ratio of blastomeres classified by the number of *Xist* domains. Each bar represents one female morula. **(H)** Immunostaining for H3K27me3 (red) in the nuclei (DAPI) of female blastocysts colabeled with CDX2 (green)-positive trophoblast cells. Rightmost panels: higher magnification of boxed regions. White arrows indicate H3K27me3 domain-negative trophoblast cells. **(I)** The ratio of trophoblast cells classified by the number of H3K27me3 domains. Each bar represents one female blastocyst. **(J)** The percentage of H3K27me3-positive and -negative trophoblast cells to total trophoblast cells in female blastocysts. Chi-square tests were used. $*p < 0.05$, $**p < 0.01$, ns, not significant. Scale bar = 25 μm.

**Fig. S2.**

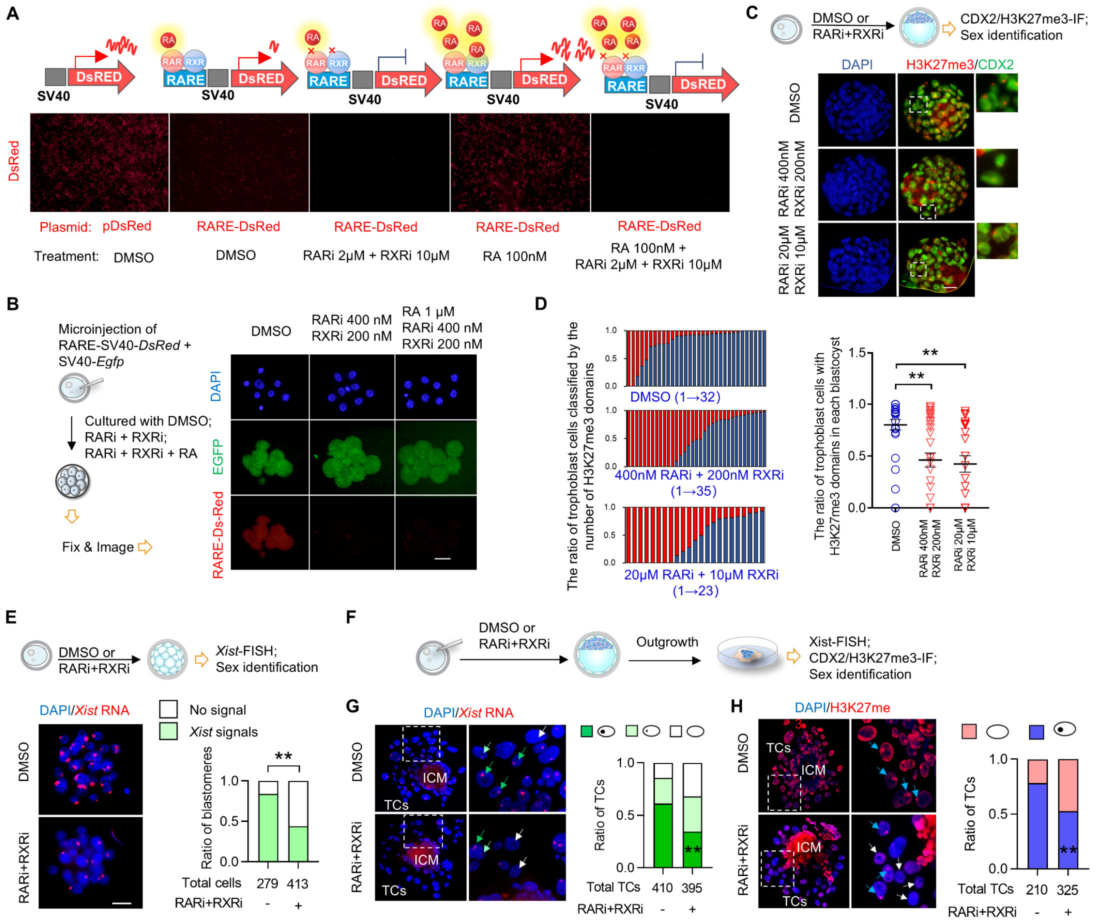


**Figure. S2. Effect of blocking nuclear retinoid receptor signaling on iXCI status. (A)** The levels of red fluorescence in HEK293T cells transfected with RA response element (RARE)-*Discosoma* red fluorescent protein (DsRed) reporter in the presence of exogenous RA or RAR/RXR pan-inhibitors, either alone or in combination. Upper panel: schematic diagram of RARE-driven reporter constructs. Results suggested the high sensitivity of RARE-DsRed reporter responding to RA signaling. **(B)** The levels of red fluorescence in 8-cell embryos. Left panel: schematic diagram of the experimental workflow, the RARE-DsRed reporter vector was injected into zygotes in the presence of exogenous RA or RAR/RXR pan-inhibitors, either alone or in combination. A SV40-EGFP vector was added as the microinjection control. Results suggested the active RA signaling in preimplantation embryos. **(C-D)** The ratio of trophoblast cells classified by the number of H3K27me3 domains in female blastocysts exposed to RAR/RXR pan-inhibitors. Upper panel: schematic diagram of the experimental workflow. Quantification of the ratio of H3K27me3-positive trophoblast cells in each blastocyst. **(E)** *Xist*-FISH analysis in female morulae exposed to RAR/RXR pan-inhibitors. Upper panel: schematic diagram of the experimental workflow. Right panel: the percentage of *Xist*-positive and -negative blastomeres to total blastomeres in female morulae. **(F)** Schematic diagram of the experimental workflow. blastocysts exposed to RAR/RXR pan-inhibitors were transferred into the standard culture medium of outgrowth, *Xist* and H3K27me3 domains were detected and quantified. **(G)** *Xist*-FISH analysis in trophoblasts of blastocyst outgrowths. Right panel: the percentage of trophoblasts without detectable *Xist* signal, with *Xist* pinpoints (pale green arrow) or with *Xist* clouds (dark green arrow). **(H)** Immunostaining for H3K27me3 in trophoblasts of blastocyst outgrowths. Right panel: the percentage of H3K27me3-positive and -negative trophoblasts. For e, g, and h, Chi-square tests were used. For others, data are mean ± s.e.m and two-tailed Student's t-tests were used to determine statistical significance. *$p < 0.05$, **$p < 0.01$, ns, not significant. Scale bar = 25 μm.

**Fig. S3.**

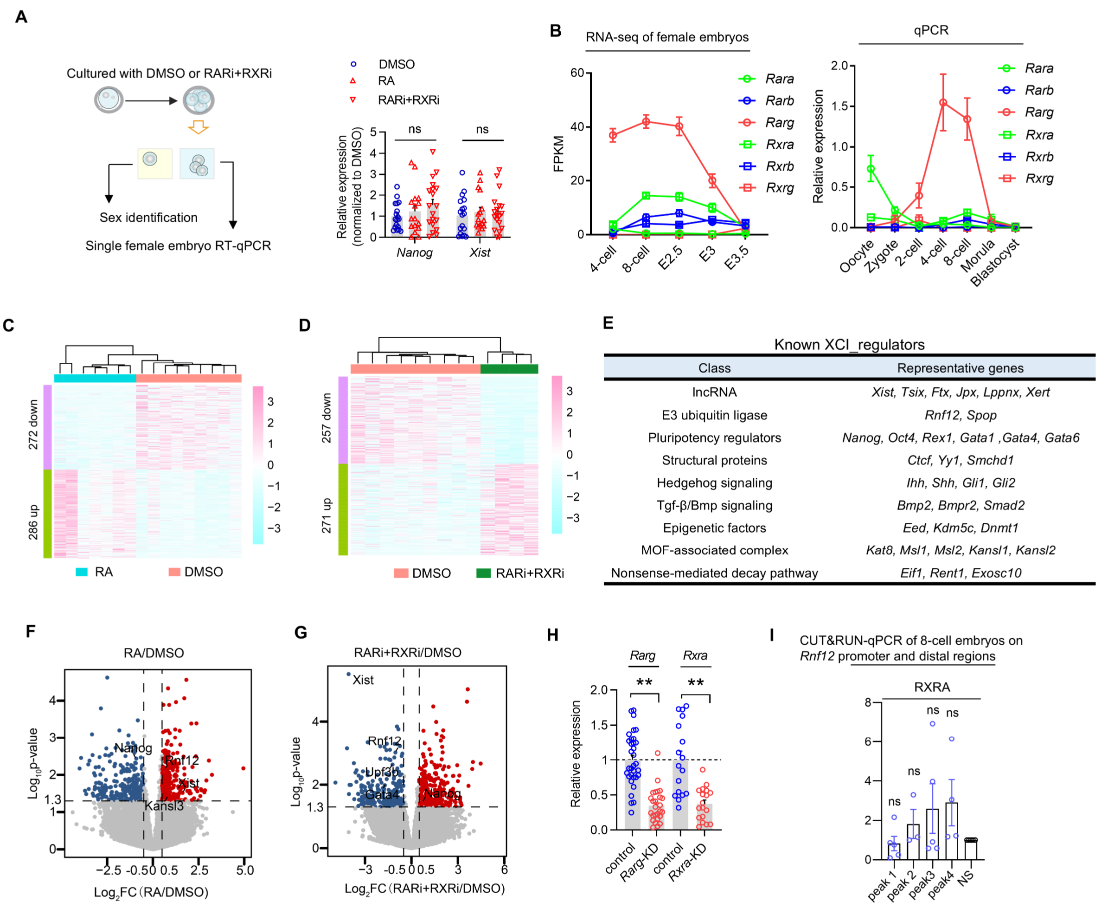


**Figure. S3. The screen of candidate mediators linking the nuclear retinoid receptor and *Xist*. (A)** Diagram illustrating embryos exposed to exogenous RA or RAR/RXR pan-inhibitors were sampled at the 4-cells for sex determination and single embryo quantitative qRT-PCR analyses. **(B)** Expression dynamics of isoforms of nuclear retinoid receptors by reanalyzing RNA-seq data GSE71442 (*3*) (left panel) and using qRT-PCR (left panel). Right paned: Relative *Xist* expression levels in female 4-cell embryos exposed to exogenous RA or RAR/RXR pan-inhibitors. **(C-D)** Heatmaps depicting expression patterns of up- and downregulated genes in response to exogenous RA (C) or RAR/RXR pan-inhibition (D). **(E)** Representative known XCI regulators and their categories. Full regulators are list in **Table. S2**. **(F-G)** Volcano plot of the significantly (red dots) up- and downregulated (blue dots) genes (FC>1.5, P<0.05). Known XCI regulators are indicated. **(H)** The detection knockdown efficiency of *Rarg* and *Rxra* in female 8-cell embryos after zygotic microinjection of *Rarg*-siRNA or *Rxra*-siRNA. **(I)** iCUT&RUN-qPCR analysis of nuclear retinoid receptors occupancy at their putative binding sites (peak1-peak4) within regulatory regions of *Rnf12* in 8-cell embryos. NS, non-specific binding site. Data are mean ± s.e.m and two-tailed Student's t-tests were used to determine statistical significance. *$p < 0.05$, **$p < 0.01$, ns, not significant. Scale bar = 25 µm.

**Fig. S4.**

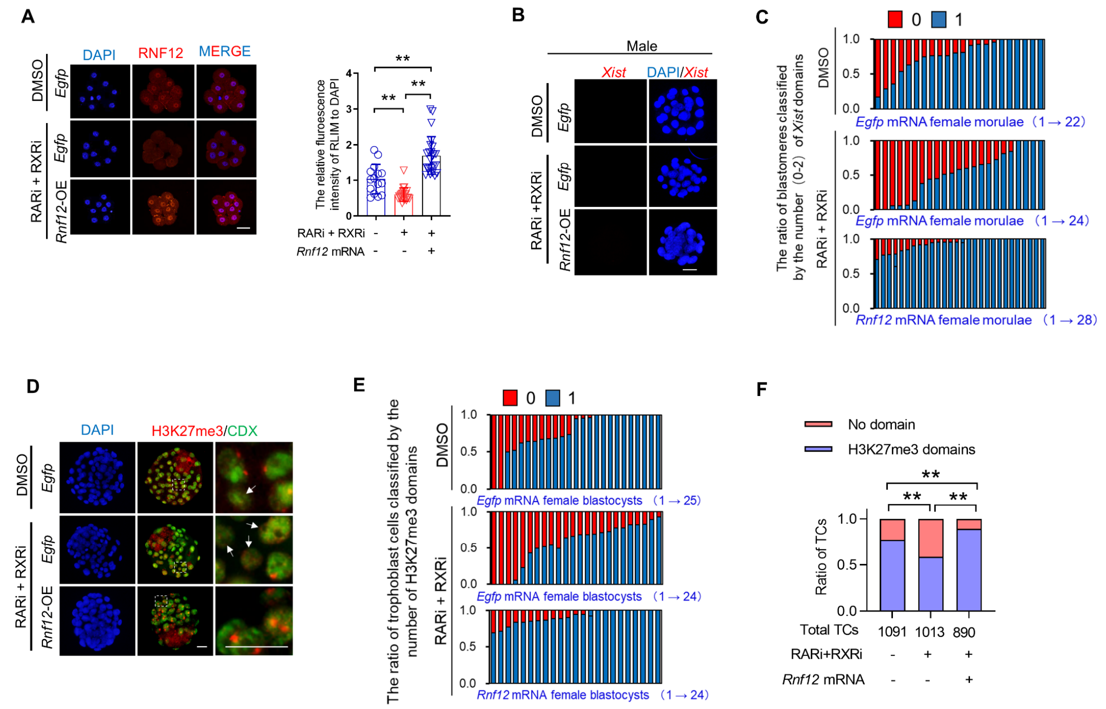


**Figure. S4. The role of *Rnf12* in mediating RA-driven iXCI. (A)** Immunostaining of RNF12 in female 8-cell embryos subjected to RAR/RXR pan-inhibition, or in combination with overexpression of *Rnf12*. Right panel: the quantification of RNF12 intensity under different conditions. **(B)** *Xist*-FISH analysis in male morulae from different intersections and maternal dietary conditions. **(C)** The ratio of blastomeres classified by the number of *Xist* domains. Each bar represents one female morula. **(D)** Immunostaining for H3K27me3 (red) in the nuclei (DAPI) of female blastocysts colabeled with CDX2 (green)-positive trophoblast cells. Rightmost panels: higher magnification of boxed regions. White arrows indicate H3K27me3 domain-negative trophoblast cells. **(E)** The ratio of trophoblast cells classified by the number of H3K27me3 domains. Each bar represents one female blastocyst. **(F)** The percentage of H3K27me3-positive and -negative trophoblast cells to total trophoblast cells in female blastocysts. For e, Chi-square tests were used. *$p < 0.05$, **$p < 0.01$, ns, not significant. Scale bar = 25 µm.

**Fig. S5.**

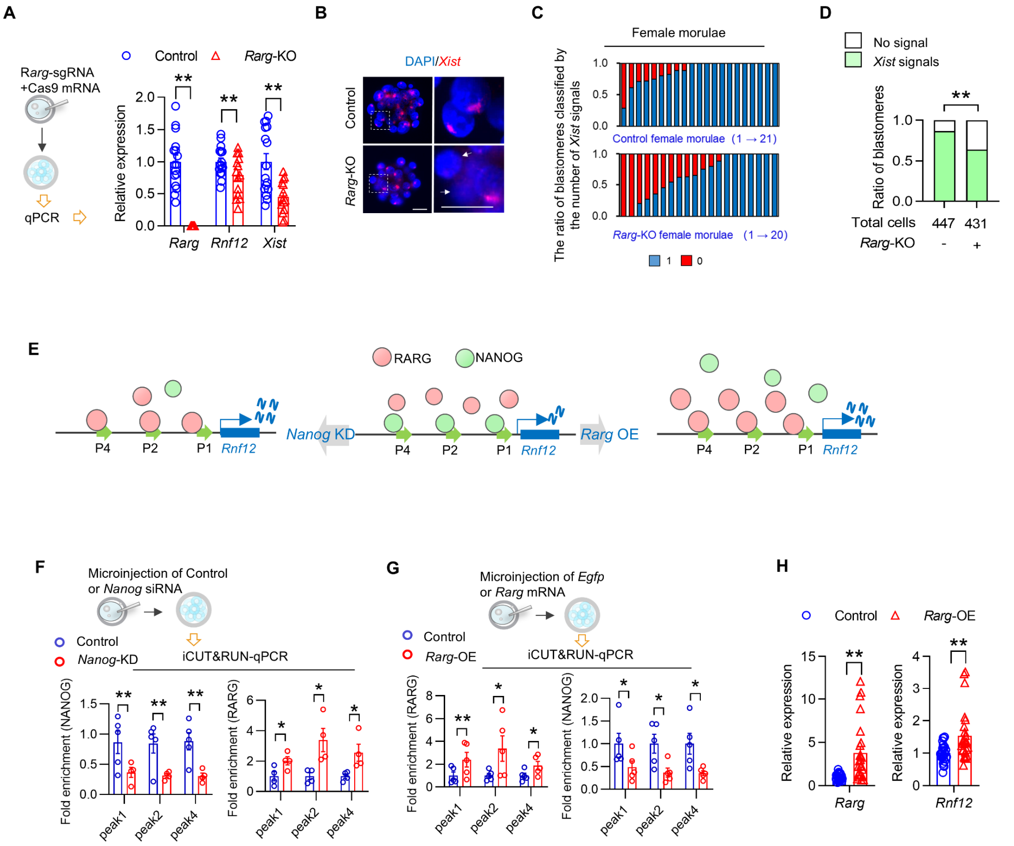


**Figure. S5. The role of *Rarg* in safeguarding iXCI. (A)** Relative expression levels of *Rarg*, *Rnf12* and *Xist* in female 8-cell embryos subjected to CRISPR/Cas9 -induced *Rarg* knockout. **(B)** *Xist*-FISH analysis in female morulae from *Rarg*-KO. White arrows indicate blastomeres without the *Xist* RNA domain. **(C)** The ratio of blastomeres classified by the number of *Xist* domains. Each bar represents one female morula. **(D)** The percentage of *Xist*-positive and -negative blastomeres to total blastomeres in female morulae. **(E)** Schematic diagram of the model in which RARG competes with NANOG for binding to peak1, peak2 and peak4 within *Rnf12* 5'region under the conditions of *Rarg* overexpression and *Nanog* knockdown respectively. **(F)** iCUT&RUN-qPCR analysis of RARG and NANOG occupancy at their putative binding sites within *Rnf12* 5' region in 8-cell embryos subjected to *Nanog* knockdown or not. **(G)** iCUT&RUN-qPCR analysis of RARG and NANOG occupancy at *Rnf12* 5' region in 8-cell embryos subjected to *Rarg* overexpression or not. **(H)** Relative expression levels of *Rarg* and *Rnf12* in female 8-cell embryos subjected to *Rarg* overexpression. For D, Chi-square tests were used. For others, data are mean ± s.e.m and two-tailed Student's t-tests were used to determine statistical significance. *$p < 0.05$, **$p < 0.01$, ns, not significant. Scale bar = 25 µm.

**Fig. S6.**

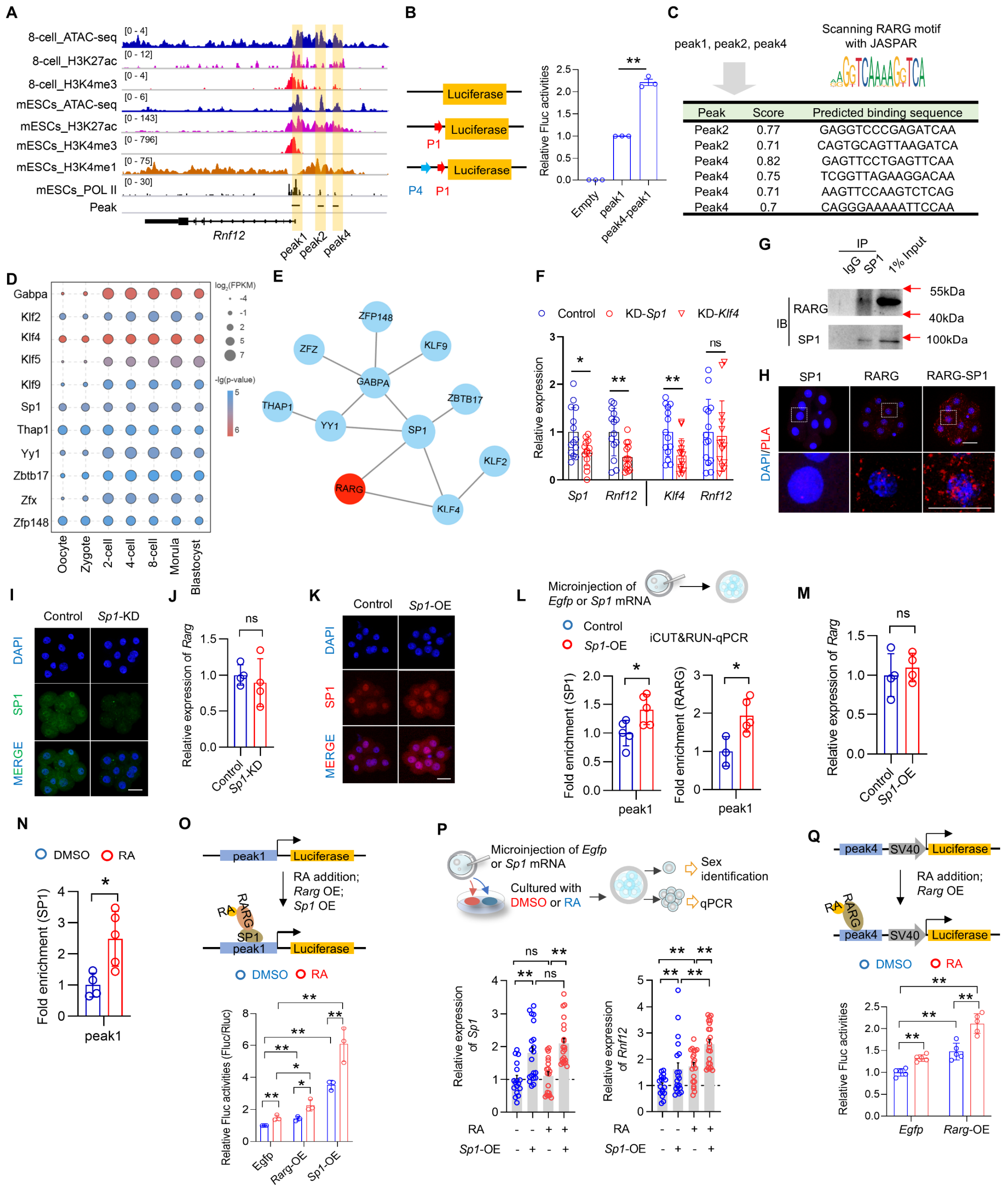


**Figure. S6. The synergistic action of SP1 and RARG in inducing *Rnf12*. (A)** Genome browser view of H3K27ac, H3K4me3, H3K4me1, RNA polymerase II ChIP-seq and ATAC-seq in 8-cell embryos and ES cells around the *Rnf12* locus. **(B)** Schematic diagram of luciferase reporter constructs containing peak1 alone or in combination with peak4. Right panel: relative activities of the presented constructions transfected into HEK293T cells. **(C)** Prediction of RARG binding sites at peak1, 2, 4 with RARG motif by JASPAR. The predicted binding sequences and potential binding scores (>0.7) are shown in the table. **(D)** Transcription factor (TF) motifs identified from peak1. Only TFs expressed at least at one stage (FPKM≥5) were included. **(E)** The interaction network among the identified TFs. **(F)** Relative expression levels of *Rnf12* in female 8-cell embryos subjected to knockdown of *Sp1* or *Klf4*. **(G)** Co-immunoprecipitation assays showing interactions of RARG with SP1 in differentiated PGK12.1 female ES cells. **(H)** Validation of the interaction between RARG and SP1 via in situ PLA in 8-cell embryos. **(I)** Immunostaining of SP1 in 8-cell embryos subjected to knockdown of *Sp1*. **(J)** Relative expression levels of *Rarg* in female 8-cell embryos subjected to knockdown of *Sp1*. **(K)** Immunostaining of SP1 in 8-cell embryos subjected to overexpression of *Sp1*. **(L)** iCUT&RUN-qPCR analysis of relative SP1 and RARG occupancy at peak1 in 8-cell embryos subjected to *Sp1* overexpression or not. **(M)** Relative expression levels of *Rarg* in female 8-cell embryos subjected to overexpression of *Sp1*. **(N)** iCUT&RUN-qPCR analysis of relative SP1 occupancy at peak1 in 8-cell embryos exposed to exogenous RA. **(O)** Relative activities of the presented constructions transfected into 293T cells exposed to RA, or in combination with *Rarg* and Sp1 overexpression. **(P)** Upper panel: schematic diagram of the experimental workflow. *Sp1* was overexpressed by microinjecting *Sp1 m*RNA into zygotes in the presence or absence of RA. Lower panel: relative expression levels of *Sp1* and *Rnf12* in female 8-cell female embryos subjected to RA supplementation or *Sp1* overexpression, either alone or in combination. (**Q**) Effect of RA supplementation, or in combination with *Rarg* overexpression, on the activity of the construction containing peak4. Data are mean ± s.e.m and two-tailed Student's t-tests were used to determine statistical significance. *$p < 0.05$, **$p < 0.01$, ns, not significant. Scale bar = 25 μm.

**Fig. S7.**

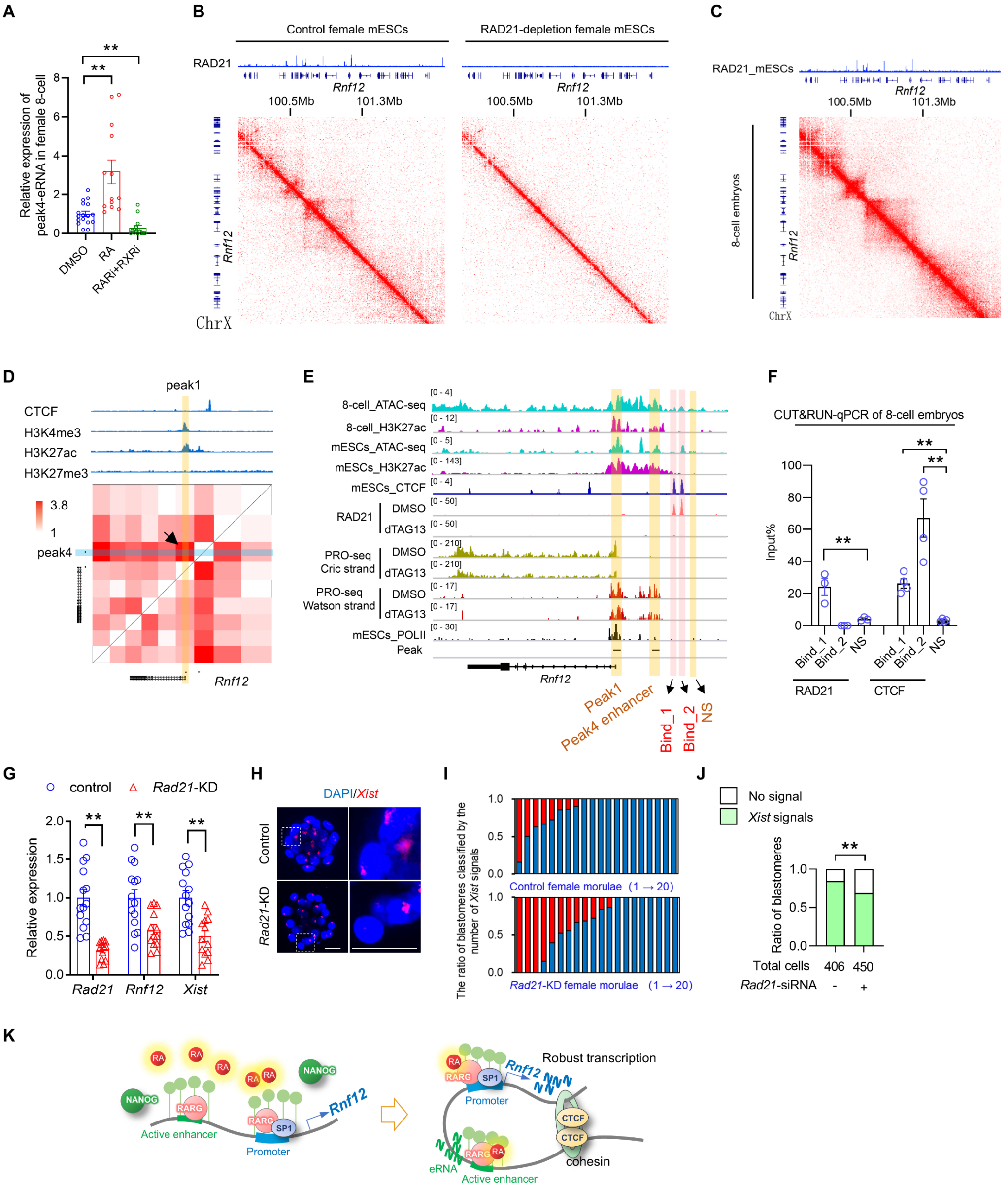


**Figure. S7. The role of enhancer-promoter loop in inducing *Rnf12*. (A)** The relative expression levels of peak4 eRNA in female 8-cell embryos exposed to exogenous RA or RAR/RXR pan-inhibitors. **(B)** Heatmap depicting Hi-C interactions for the X chromosome in wild-type and *Rad21* depletion female ES cells (*4*). Results showed that peak1 and peak4 located in the same topologically associating domain (TAD), whose formation was mediated by RAD21. **(C)** Heatmap depicting Hi-C interactions for the X chromosome in 8-cell embryos (*5*), which showed that peak1 and peak4 located in the same TAD. **(D)** Heatmap depicting high-resolution Hi-C interactions around the *Rnf12* locus. The site of peak1 enriched with H3K27ac and H3K4me3 is highlighted with yellow. Peak4 site is highlighted with blue. Results indicate the high interaction frequency of peak1 and peak4. **(E)** Genome browser view of H3K27ac, RNA polymerase II ChIP-seq and ATAC-seq, as well as RAD21-depletion (showed by dTAG13 treatment) PRO-seq data, around the *Rnf12* locus in 8-cell embryos and ES cells. The peak1, peak4, and putative binding sites of CTCF (Bind_1 and Bind_2), are highlighted. **(F)** iCUT&RUN-qPCR analysis of RAD21 and CTCF occupancy at *Rnf12* upstream in 8-cell embryos. **(G)** Relative expression levels of *Rnf12* and *Xist* in female 8-cell embryos. **(H)** *Xist*-FISH analysis in female morulae subjected to knockdown of *Rad21.* **(I)** The ratio of blastomeres classified by the number of *Xist* domains. Each bar represents one female morula. **(J)** The percentage of *Xist*-positive and -negative blastomeres to total blastomeres in female morulae. **(K)** Schematic diagram of the model in which a RAD21/CTCF-anchored E-P loop structure forms and is essential for inducing Rnf12 expression. For A, F, and G, data are mean ± s.e.m and two-tailed Student's t-tests were used to determine statistical significance. For J, Chi-square tests were used. *$p < 0.05$, **$p < 0.01$, ns, not significant. Scale bar = 25 μm.

**Fig. S8.**

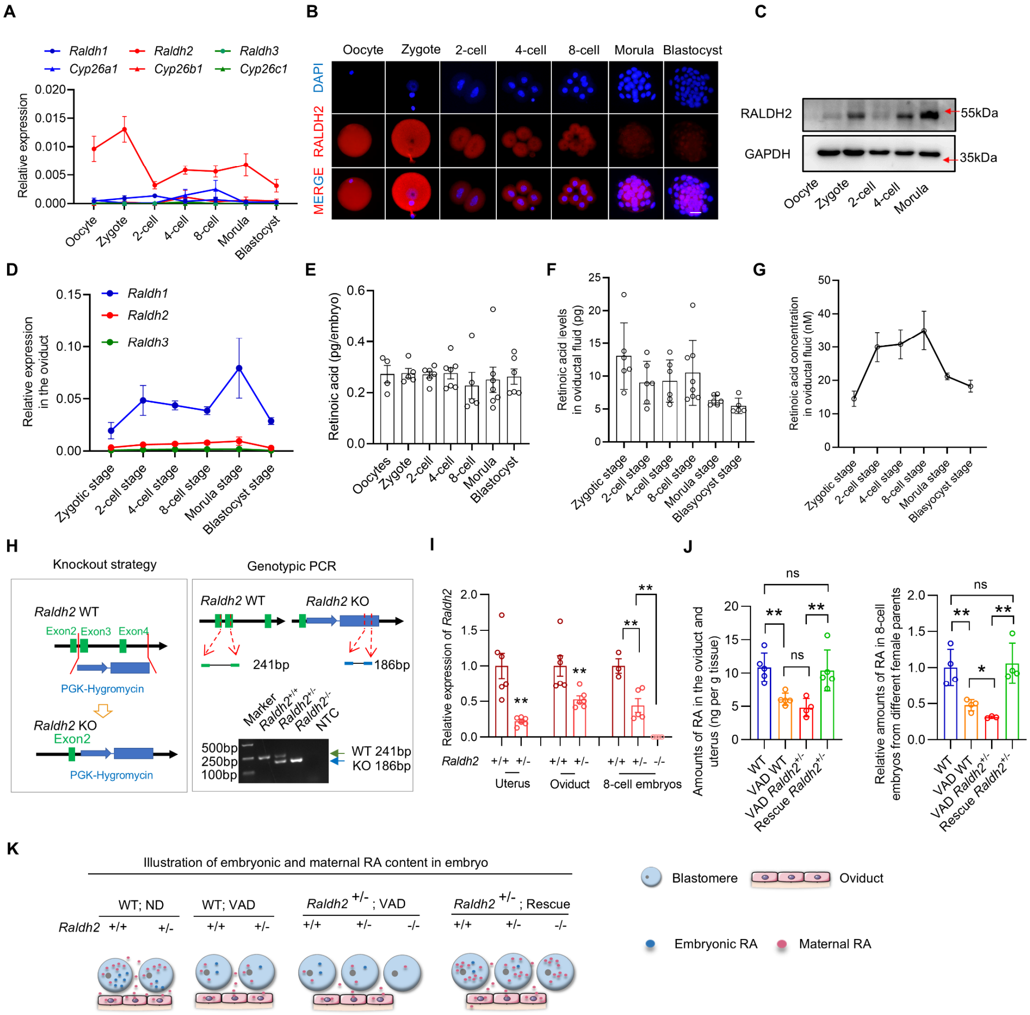


**Figure. S8. The expression patterns of RA-synthesizing enzymes and the amounts of RA during preimplantation development. (A)** The expression dynamics of RA-synthesizing and -degrading enzymes in preimplantation embryos. **(B-C)** Representative immunostaining (B) and Western blot analyses (C) for RALDH2 in preimplantation embryos. **(D)** The expression dynamics of RA-synthesizing enzymes in the oviduct during the preimplantation stages. **(E)** Detection of RA amounts in embryos at different preimplantation stages by ELISA. **(F)** Detection of RA amounts in oviductal fluid during the preimplantation stages by ELISA. **(G)** The dynamics of RA concentration in oviductal fluid during the preimplantation stages showed high RA concentration in oviduct during iXCI initiation. **(H)** A schematic diagram of the previously targeted *Raldh2* allele with a deletion of exon 3 and 4 (*6*), and genotypic detection of wild-type, *Raldh2*$^{+/-}$, *Raldh2*$^{-/-}$ embryos. Primers used for genotyping and expected band sizes are shown in the upper panel. **(I)** Relative expression levels of *Raldh2* in the uterus and oviduct of the wild-type and *Raldh2*$^{+/-}$ female parents, as well as in their resulting wild-type, *Raldh2*$^{+/-}$, *Raldh2*$^{-/-}$ embryos. **(J)** Detection of RA amounts in the oviduct and uterus (ng per g tissue) of female mice with different genotypes and dietary conditions by LC-MS/MS. Right panel: Detection of the relative RA amounts in 8-cell embryos from female parents with different genotypes and dietary conditions by ELISA. **(K)** A summary of source and supply of RA under different parental genotypes and maternal dietary conditions. Data are mean ± s.e.m and two-tailed Student’s t-tests were used to determine statistical significance. *$p < 0.05$, **$p < 0.01$, ns, not significant. Scale bar = 25 μm.

**Fig. S9.**

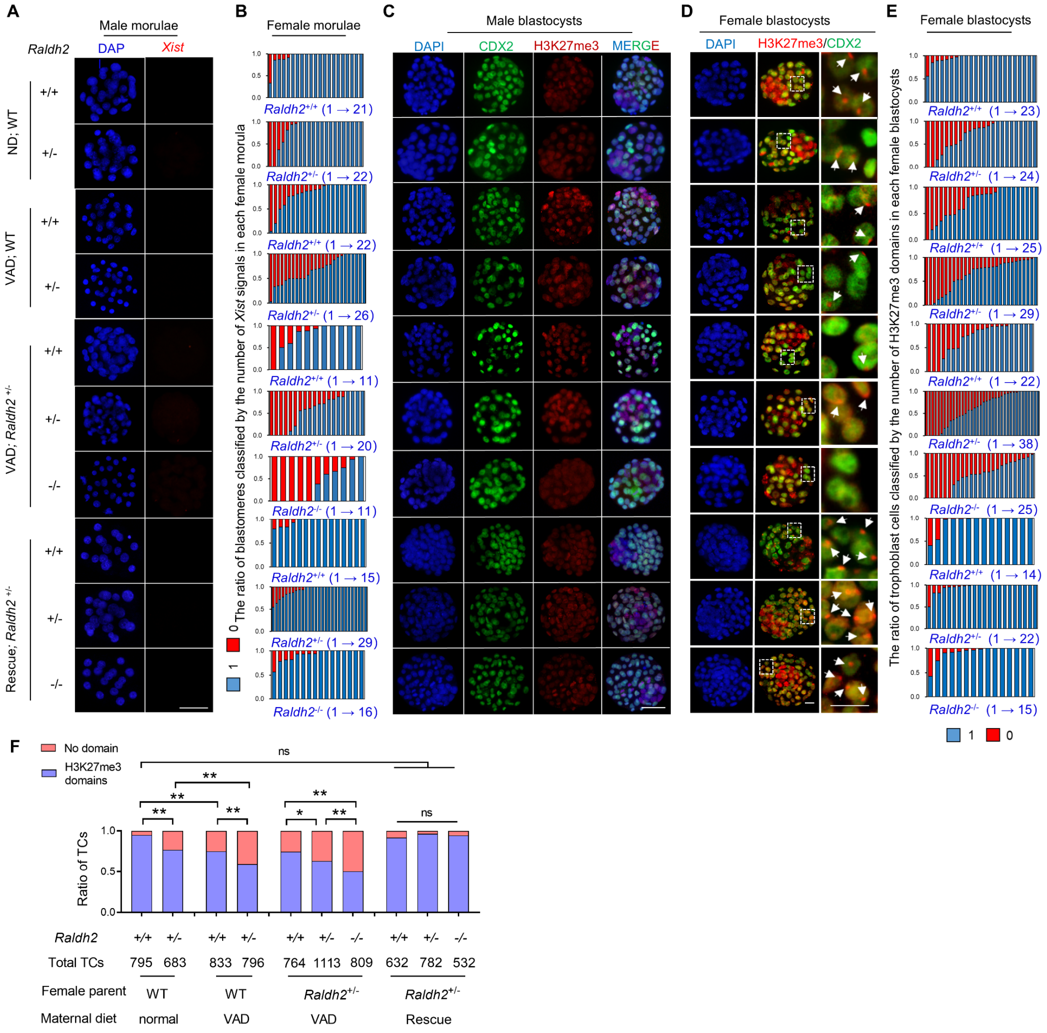


**Figure. S9. RA deficiency impairs iXCI in preimplantation female embryos. (A)** *Xist*-FISH analysis in male morulae from different intersections and maternal dietary conditions. **(B)** The ratio of blastomeres classified by the number of *Xist* domains present in Figure 1g. Each bar represents one female morula. **(C-D)** Representative immunostaining for H3K27me3 (red) in the nuclei (DAPI) of male (C) and female (D) blastocysts colabeled with CDX2 (green)-positive trophoblast cells. Rightmost panels in d: higher magnification of boxed regions. White arrows indicate CDX2 positive trophoblast cells with the H3K27me3 domain. **(E)** The ratio of trophoblast cells classified by the number of H3K27me3 domains in d. Each bar represents one female blastocyst. **(F)** The percentage of H3K27me3-positive and -negative trophoblast cells to total trophoblast cells in female blastocysts. Chi-square tests were used to determine statistical significance. $*p < 0.05$, $**p < 0.01$, ns, not significant. Scale bar = 25 µm.

**Fig. S10.**

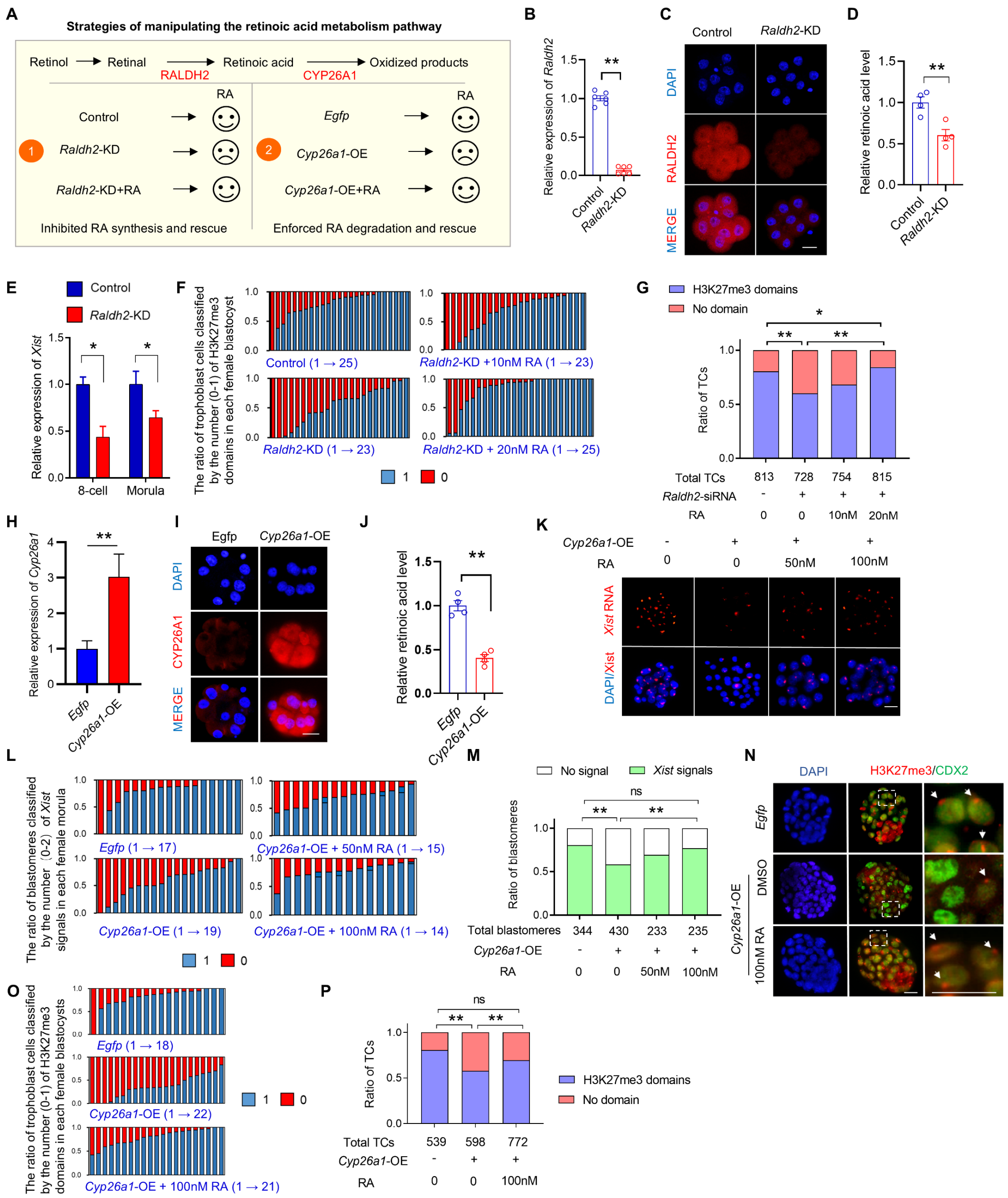


**Figure. S10. Effect of modulating embryonic RA signaling on iXCI status. (A)** Schematic diagram of two strategies for modulating embryonic RA signaling by microinjecting *Raldh2* siRNA and *Cyp26a1* mRNA, in the presence or absence of exogenous RA. Upper panel: the synthesis and degradation pathway of RA. **(B-C)** Relative expression levels of *Raldh2* (B) and immunostaining of RALDH2 (C) in 8-cell embryos injected with *Raldh2* -siRNA and control siRNA. **(D)** Detection of the relative RA level in 8-cell embryos after knockdown of *Raldh2* by ELISA. **(E)** Relative expression levels of *Xist* in female 8-cell embrys and morulae after knockdown of *Raldh2*. **(F)** The ratio of trophoblast cells classified by the number of H3K27me3 domains in blastocysts subjected to *Raldh2* knockdown alone, or in combination with exogenous RA supplementation. Each bar represents one female blastocyst. **(G)** The percentage of

H3K27me3-positive and -negative trophoblast cells to total trophoblast cells in female blastocysts. **(H-I)** Relative expression levels of *Cyp26a1* (h) and immunostaining of CYP26A1 (i) in 8-cell embryos injected with *Cyp26a1 mRNA* and *Egfp* mRNA (control). **(J)** Detection of the relative RA level in 8-cell embryos overexpressing *Raldh2* by ELISA. **(K)** *Xist*-FISH analysis in female morulae subjected to *Cyp26a1* overexpression alone, or in combination with exogenous RA supplementation. **(L)** The ratio of blastomeres classified by the number of *Xist* domains present in k. Each bar represents one female morula. **(M)** The percentage of *Xist*-positive and -negative blastomeres to total blastomeres in female morulae. **(N)** Representative immunostaining for H3K27me3 (red) in the nuclei (DAPI) of female blastocysts subjected *Cyp26a1* overexpression alone, or in combination with exogenous RA supplementation. Rightmost panels: higher magnification of boxed regions. White arrowheads indicate CDX2 positive trophoblast cells with the H3K27me3 domain. **(O)** The ratio of trophoblast cells classified by the number of H3K27me3 domains in female blastocysts. **(P)** The percentage of H3K27me3-positive and -negative trophoblast cells to total trophoblast cells in female blastocysts. For B, D, E, H, and J, data are mean ± s.e.m and two-tailed Student's t-tests were used to determine statistical significance. For G, M, and P, Chi-square tests were used. *$p < 0.05$, **$p < 0.01$, ns, not significant. Scale bar = 25 μm.

#### Fig. S11.

**A**

| Group | Female | IS | E7.5 | IS / female | E7.5 / female | Resorption rate |
|---|---|---|---|---|---|---|
| Normal diet; WT ♀ X WT ♂ | 12 | 135 | 132 | 11.25 | 11.00 | 2% |
| Normal diet; WT ♀ X *Raldh2*+/- ♂ | 41 | 371 | 312 | 9.05 | 7.61 | 16% (3.3E-05) |
| VAD; WT♀ X *Raldh2*+/- ♂ | 39 | 252 | 193 | 6.46 | 4.95 | 23% (6.1E-08) |
| VAD; *Raldh2*+/- ♀ X *Raldh2*+/- ♂ | 62 | 380 | 278 | 6.13 | 4.48 | 27% (1.1E-09) |
| Rescue diet; *Raldh2*+/- ♀ X *Raldh2*+/- ♂ | 20 | 227 | 210 | 11.35 | 10.50 | 7% (0.03) |

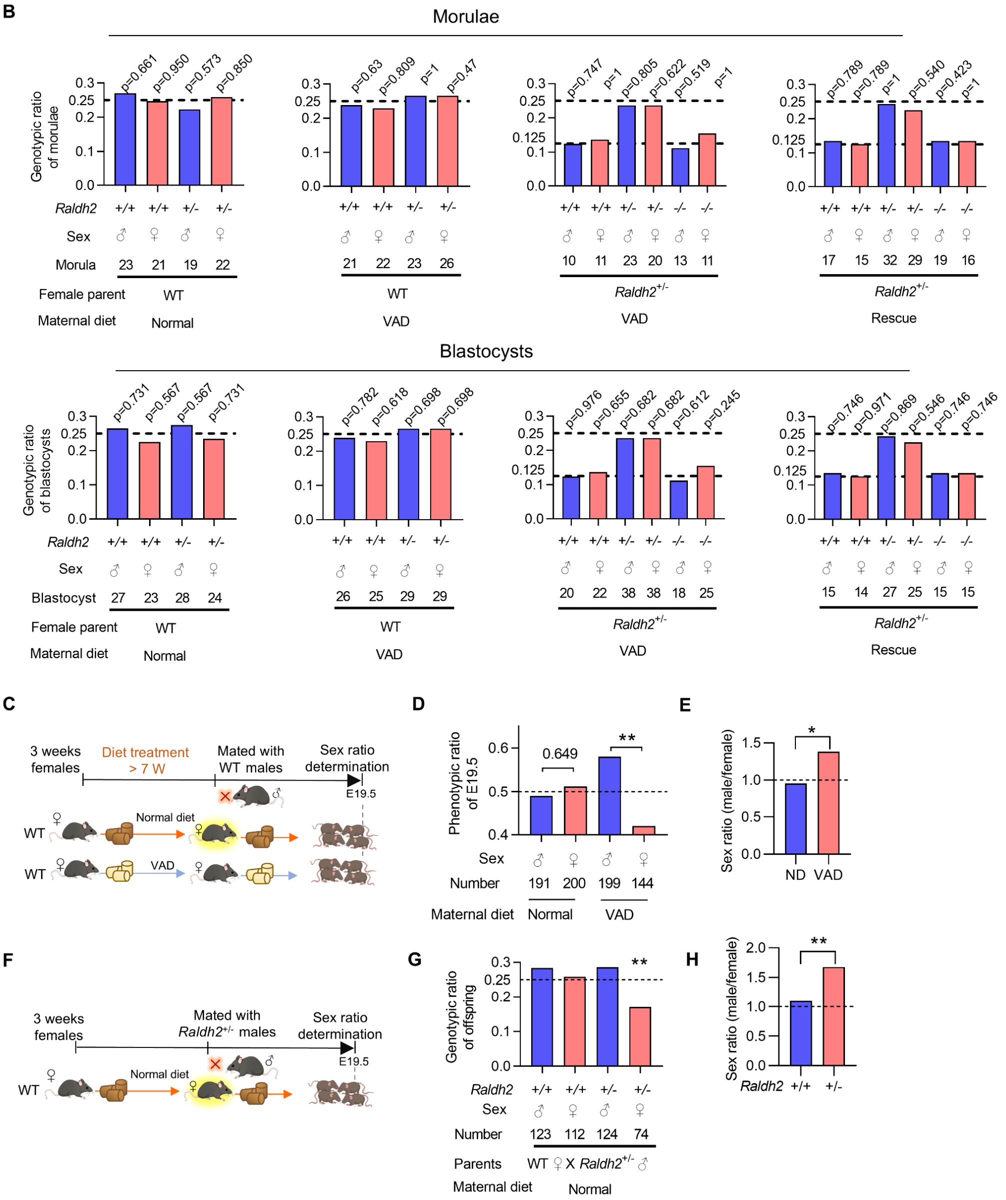


**Figure. S11. The effect of maternal RA deficiency and embryonic RA synthetic defects on embryogenesis. (A)** The developmental rate of E7.5 embryos from different crosses and dietary conditions, showing the RA deficiency-associated embryonic defects and rescued embryogenesis by maternal RA supplementation. **(B)** Genotype data of morulae (H) and blastocysts (I) from female mice with different genotypes and dietary conditions. The dashed line shows the expected

Mendelian ratio and the underlined number shows the observed number of embryos. **(C)** Schematic diagram of the experimental design. 3 weeks wild-type (WT) female mice were fed a normal diet (ND) or a VA deficiency diet (VAD) for over 7 weeks. Afterward, the ND and VAD female mice were mated with WT males. E19.5 pups from the ND and VAD pregnant mice were used to calculate the sex ratio. **(D)** Phenotypic ratio of E19.5 pups exposed to maternal vitamin A-deficient (VAD) diet or normal diet. **(E)** Sex ratio of offspring exposed to maternal vitamin A-deficient (VAD) diet or normal diet (ND). **(F)** Schematic diagram of the experimental design. **(G)** Phenotypic data of offspring from the female WT (normal diet) and male *Raldh2*$^{+/-}$ cross. **(H)** Sex ratio of WT and *Raldh2*$^{+/-}$ offspring from female WT (normal diet) and male *Raldh2*$^{+/-}$ cross. Chi-square tests were used. *$p < 0.05$, **$p < 0.01$, ns, not significant. Scale bar = 25 μm.

**Fig. S12.**

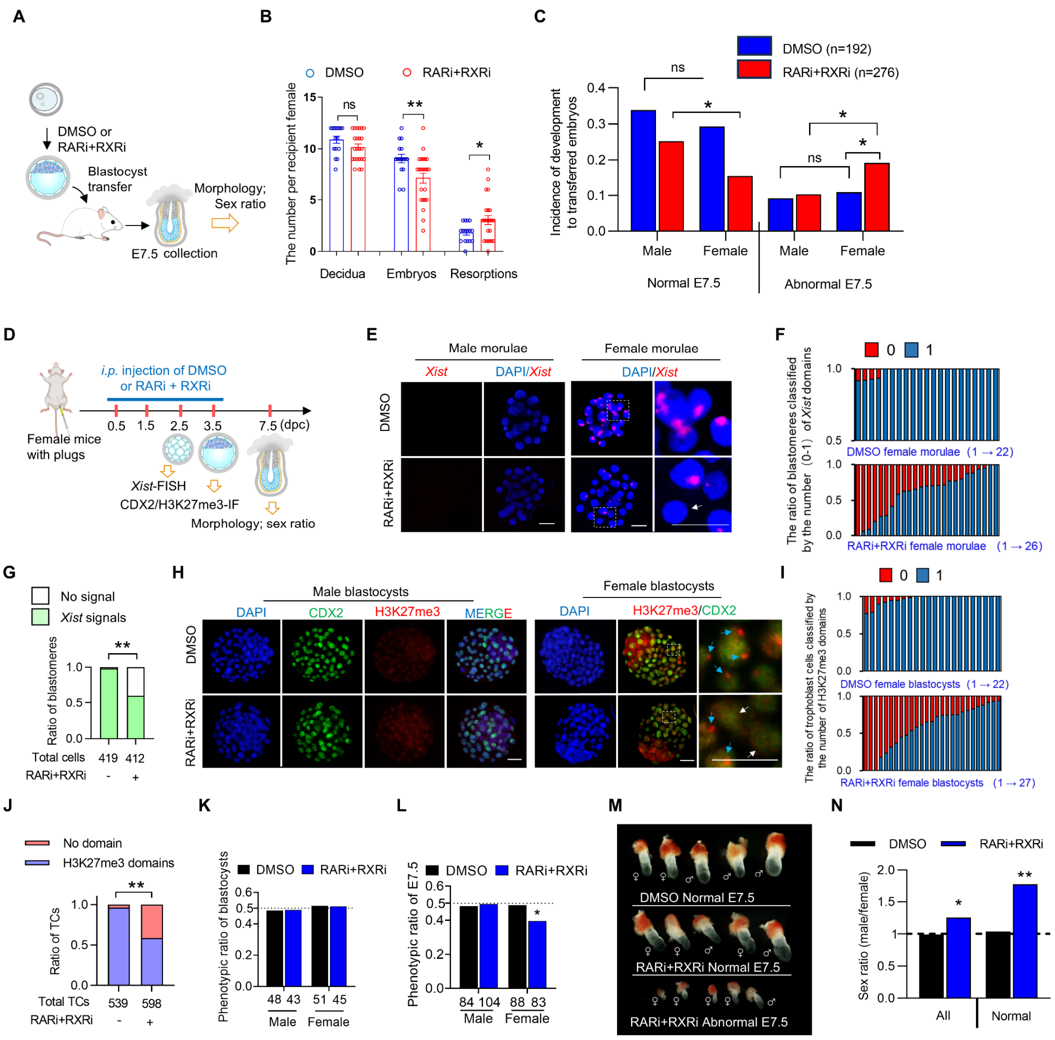


**Figure. S12. Effect of in vivo blockage of nuclear retinoid receptor signaling on iXCI status and female embryogenesis. (A)** Schematic diagram of the experimental workflow. Blastocysts exposed to RAR/RXR pan-inhibitors were transferred into normal recipients. **(B)** The number of the decidua, recovered embryos and resorptions per recipient at E7.5. **(C)** Incidence of development of male and female embryos with different morphologies at E7.5 after embryo transfer. **(D)** Schematic diagram of the experimental workflow. RAR/RXR pan-inhibitors were intraperitoneally injected during the preimplantation window of iXCI onset. iXCI status and female embryogenesis were detected at indicated time points. Results suggested that the blockage of RA signaling *in vivo* impaired *Xist* RNA coating and H3K27me3 domains, and further led female-biased defects in post-implantation embryos. **(E)** *Xist*-FISH analysis in male and female morulae. White arrows indicate *Xist*-negative blastomeres. **(F)** The ratio of blastomeres classified by the number of *Xist* domains. Each bar represents one female morula. **(G)** The percentage of *Xist*-positive and -negative blastomeres (with arrow in c) to total blastomeres in female morulae. **(H)** Immunostaining for H3K27me3 (red) in the nuclei (DAPI) of male and female blastocysts colabeled with CDX2 (green)-positive trophoblast cells. Rightmost panels in b and e: higher magnification of boxed regions. Blue arrows indicate

H3K27me3 domain-positive trophoblast cells. White arrows indicate H3K27me3-negative trophoblast cells. **(I)** The ratio of trophoblast cells classified by the number of H3K27me3 domains. Each bar represents one female blastocyst. **(J)** The percentage of H3K27me3-positive and -negative trophoblast cells to total trophoblast cells in female blastocysts. **(K-L)** Phenotypic ratio of male and female blastocysts (k) and E7.5 embryos (l) subjected to intraperitoneal RAR/RXR pan-inhibition. **(M)** Representative images of E7.5 embryos with normal or abnormal morphologies. **(N)** Sex ratios of all recovered E7.5 embryos or those with normal morphology. For c, g, j, k, l, and m, Chi-square tests were used to determine statistical significance. $*p < 0.05$, $**p < 0.01$, ns, not significant. Scale bar = 25 µm.


**Table S1. (separate file)**

PCR and qPCR primers; siRNA and sgRNA targeted sequences.

**Table S2. (separate file)**

The summary of known XCI regulators.

**Table S3. (separate file)**

Gene expression (TPM) of female 8-cell embryos after DMSO, RA, or RA receptors inhibitors treatment.
